# Pharmacological Inhibition of Epac1 Protects against Pulmonary Fibrosis by Blocking FoxO3a Neddylation

**DOI:** 10.1101/2024.09.13.612935

**Authors:** Katherine Jankowski, Sarah E Lemay, Daniel Lozano-ojalvo, Leticia Perez Rodriguez, Mélanie Sauvaget, Sandra Breuils-Bonnet, Karina Formoso, Vineeta Jagana, Shihong Zhang, Javier Milara, Julio Cortijo, Irene C. Turnbull, Steeve Provencher, Sebastien Bonnet, Jordi Orchando, Frank Lezoualc’h, Malik Bisserier, Lahouaria Hadri

**Affiliations:** Center of Excellence for Translational Medicine and Pharmacology/Department of Pharmacological Sciences, Icahn School of Medicine, Mount Sinai, New York, NY 10029, USA; Cardiovascular Research Institute, Icahn School of Medicine at Mount Sinai, New York, NY 10029, USA; Institut Universitaire de Cardiologie et de Pneumologie de Québec Research Center, Université Laval, Quebec City, QC, Canada; Pulmonary Hypertension Research Group, Université Laval, Quebec City, QC, Canada; Department of Medicine, Université Laval, Quebec City, QC, Canada; Oncological Sciences, Icahn School of Medicine at Mount Sinai, New York, NY 10029, USA; Institut des Maladies Métaboliques et Cardiovasculaires, Inserm, Université de Toulouse III-Paul Sabatier, France; Department of Cell Biology and Anatomy and Physiology, New York Medical College, Valhalla, NY 10595, USA; Department of Pharmacology, Faculty of Medicine, University of Valencia, Valencia, Spain. Biomedical Research Networking Center on Respiratory Diseases (CIBERES), Health Institute Carlos III, Madrid, Spain

**Keywords:** Epac1, pulmonary fibrosis, profibrotic markers, neddylation

## Abstract

**Background:** Idiopathic Pulmonary fibrosis (IPF) is characterized by progressive scarring and fibrosis within the lungs. There is currently no cure for IPF; therefore, there is an urgent need to identify novel therapeutic targets that can prevent the progression of IPF. Compelling evidence indicates that the second messenger, cyclic adenosine monophosphate (cAMP), inhibits lung fibroblast proliferation and differentiation through the classical PKA pathway. However, the contribution of the <u>e</u>xchange <u>p</u>rotein directly <u>a</u>ctivated by <u>c</u>AMP 1 (Epac1) to IPF pathophysiological processes is yet to be investigated.

**Objective:** To determine the role of the cAMP-binding protein Epac1 in the progression of IPF.

**Methods:** We used lung samples from IPF patients or healthy controls, mouse lung samples, or lung fibroblast isolated from a preclinical mouse model of PF induced by bleomycin intratracheal injection. The effect of bleomycin (BLM) treatment was determined in Epac1 knock-out mice or wild-type littermates. Epac1 expression was modulated *in vitro* by using lentiviral vectors or adenoviruses. The therapeutic potential of the Epac1-selective pharmacological inhibitor, AM-001, was tested *in vivo* and *in vitro,* using a bleomycin mouse model of PF and an *ex vivo* precision-cut lung slices (PCLs) model of human lung fibrosis.

**Results:** Epac1 expression was increased in the lung tissue of IPF patients, in IPF-diseased fibroblasts and in BLM-challenged mice. Furthermore, Epac1 genetic or pharmacological inhibition with AM-001 decreased normal and IPF fibroblast proliferation and the expression of profibrotic markers, αSMA, TGF-β/SMAD2/3, and interleukin-6 (IL-6)/STAT3 signaling pathways. Consistently, blocking Epac1 protected against BLM-induced lung injury and fibrosis, suggesting a therapeutic effect of Epac1 inhibition on PF pathogenesis and progression. Global gene expression profiling revealed a decrease in the key components of the profibrotic gene signature and neddylation pathway in Epac1-deficient lung fibroblasts and IPF human-derived PLCs. Mechanistically, the protective effect of Epac1 inhibition against PF development involves the inhibition of FoxO3a neddylation and its subsequent degradation by NEDD8, and in part, by limiting the proliferative capacity of lung-infiltrating monocytes.

**Conclusions:** We demonstrated that Epac1 is an important regulator of the pathological state of fibroblasts in PF and that small molecules targeting Epac1 can serve as novel therapeutic drugs against PF.

## INTRODUCTION

Idiopathic pulmonary fibrosis (IPF) is a severe and progressive age-related disease that results in respiratory failure and, eventually, death^1–5^. Despite extensive research efforts in experimental and clinical studies, the incidence of IPF continues to rise with increasing morbidity, with a median survival of 3.8 years^6^. To date, only two therapeutic options have emerged, which slow disease progression and do not reverse the existing fibrosis. Current paradigms of pulmonary fibrosis (PF) include prominent events in both injury and repair^7^, where chronic or unresolved injury initiates a tissue response characterized by the recruitment of inflammatory cells, concomitant activation and proliferation of fibroblasts to profibrogenic myofibroblasts, and exaggerated extracellular matrix (ECM) deposition that distorts the pulmonary architecture and compromises lung function^8^. TGF-β/SMAD3 signaling is considered a “master switch” in the regulation of fibrosis^9–12^ and predisposes elderly people to the development of IPF^13–17^. We and others have reported that counteracting TGF-β signaling prevents and attenuates PF^18–20^. Increasing evidence suggests that a positive feedback loop between interleukine-6 (IL-6) and TGF-β may be associated with PF^20–22^. Owing to the complexity and multicellular pathophysiology of PF, alternative pharmacological therapies to reverse this disease are needed.

Cyclic adenosine monophosphate (cAMP) is an important second messenger that regulates many physiological and pathophysiological processes via two main direct effectors, protein kinase A (PKA) and exchange proteins directly activated by cAMP, Epac1 and Epac2^23–28^. For many years, PKA has been described as the “classic” effector responsible for diverse cAMP-mediated functions. With respect to fibrosis formation, various studies have shown that cAMP inhibits fibroblast activation through the PKA pathway ^29–31^. Given the emerging role of Epac1 in modulating inflammatory and fibrotic responses in other organ systems, its investigation in the context of lung fibrosis offers a promising avenue for identifying new therapeutic targets ^32^. Evidence that Epac1 and TGF-β1 trigger acute AKT phosphorylation suggests that the phosphoinositide 3-kinase (PI3K) axis may regulate TGF-β receptor signaling in lung fibrosis ^33^. FoxO3a-deficient mice exhibit enhanced susceptibility to bleomycin (BLM) and increased PF and mortality^34^. However, the specific mechanisms by which Epac1 may contribute to IPF pathogenesis, including its potential role in mediating FB proliferation and differentiation beyond the classical PKA pathway, remain unexplored. Addressing these knowledge gaps by elucidating the role of Epac1 in IPF pathogenesis and progression could reveal novel therapeutic avenues, directly responding to the urgent need for more effective IPF treatments. The specific mechanisms by which Epac1 may contribute to IPF pathogenesis, including its potential role in mediating fibroblast proliferation and differentiation beyond the classical PKA pathway, remain unexplored. Addressing these knowledge gaps by elucidating the role of Epac1 in IPF pathogenesis and progression could reveal novel therapeutic avenues, directly responding to the urgent need for more effective IPF treatments.

In the present study, we investigated the role of Epac1 and its mechanism of action in PF progression using a combination of genetic and pharmacological approaches. We found that the inhibition of Epac1 *in vitro* and *in vivo* prevented the growth of IPF fibroblasts and PF development, respectively. Mechanistically, Epac1 blockade prevented TGF-β-induced SMAD2/3 and IL-6/STAT3 signaling pathways. Of particular importance, pharmacological inhibition of Epac1 with the small molecule AM-001 restored the expression of a key regulator of fibroblast activation, FoxO3a by preventing ubiquitin-mediated proteasomal degradation via inhibition of the neddylation pathway.

## MATERIALS AND METHODS

### Patient and Sample Collection

In this study, RNA, protein, and formalin-fixed paraffin-embedded tissue sections were obtained from human IPF patients and healthy control donors. Human lung tissue was procured from patients with IPF (*n*=8) who underwent surgery as part of an organ transplantation program, and the lung explant samples from healthy control donors were obtained from the Organ Transplant Program of the University General Consortium Hospital of Valencia (*n*=8). All samples were archived and anonymized (**Supplementary Tables 1 and 2**). The study protocol was approved by the local research and independent ethics committee of the University General Consortium Hospital of Valencia (CEIC/2013). Written informed consent was obtained from each participant before inclusion in the study.

### *Ex vivo* model of human precision-cut lung slices

The use of human lung tissues for precision-cut lung slices was approved by the Biosafety and Ethics Committees of Laval University and the Institut Universitaire de Cardiologie et de Pneumologie de Québec (IUCPQ) (CER #22390). Human lung biopsies were obtained from patients with pulmonary fibrosis. The tissues were perfused with 2% (m/V) low-melting agarose (Sigma-Aldrich) through the visible vessels and bronchi. The tissues were then sectioned into 400 µm thick slices using a Compresstome® VF-310-0Z Vibrating Microtome (Precisionary, USA).

These lung slices were sequentially placed individually in a 24-well plate containing 1 mL of medium and allowed two days to recover before treatment initiation. The slices were cultured for ten days in DMEM supplemented with 10% fetal bovine serum (FBS), 100 units/mL penicillin, 0.1 mg/mL streptomycin, 0.25 mg/mL amphotericin B, and 0.03 mg/mL Gentamicin, along with AM-001 (20 µM) or its vehicle (DMSO). The medium was replaced every two days. On the tenth day of treatment, the lung slices were fixed in formaldehyde, embedded in paraffin, and processed for Masson’s trichrome staining and immunofluorescent labeling or snap-frozen to assess variations in protein expression by western blotting. The clinical information of the patients included in this study is presented in **Supplementary Table 3**.

### Reagents

The non-hydrolyzable Epac1 preferential agonist, Sp-8-(4-chlorophenylthio)-2′-O-methyl-cAMP (Sp-8-pCPT), was purchased from BioLog (Cat# C052). This compound is abbreviated here as 8-CPT (also known as 8-pCPT-cAMP). The Epac1 inhibitor AM-001 was synthesized using previously published methods ^35^. Pevonedistat (MLN4924) is a potent and selective inhibitor of the NEDD8-activating enzyme (NAE) with an IC50 of 4.7 nM, obtained from MedChem Express (Cat# HY-70062). All the reagents are listed in **Supplementary Table 4**.

### Generation and Use of Epac1-Deficient Mice (Epac1^-/-^)

Epac1-deficient mouse line (Epac1^-/-^) has been previously described ^36^. Epac1 knock-out mice were generated by Genoway through the insertion of loxP sequences within introns 7 and 15 of RAPGEF3. Desmin-Cre (C57BL/6 background) transgenic females, which sporadically express Cre recombinase in oocytes, were crossed with Epac1^flox/flox^ (C57BL/6-SV129 background) males to generate Desmin-Cre-Epac1-/- and Epac1^-/-^ mice. Genotyping of these mice was performed by PCR using the following primers: 5′-GTTTGCCTGCCTGAATGTCT-3’, 5′-ATCTTGCCCTTCCCAGAAGT-3’, and 5′-CATGAAGCAAAGACAGTTGACATC-3’. In this study, 12-week-old mice (C57BL/6 or C57BL/6-SV129 background for Epac1 knock-out and control littermates) were used. All animal procedures were conducted according to the Institutional Animal Experimentation Guidelines and were approved by the French Ministry of Agriculture. Additionally, our study adhered to the Guide for the Care and Use of Laboratory Animals, as outlined in Directive 2010/63/EU of the European Parliament. The mice were housed in a pathogen-free facility, and all animal experiments were approved by the Animal Care and Use Committees of the University of Toulouse.

### Cell Culture

Normal human lung fibroblasts (NHL-FBs; Cat# CC-2512) and disease-related human lung fibroblasts (IPF-FBs; Cat# CC-7231) were procured from Lonza Inc. (Allendale, NJ, USA) and cultured according to recommended guidelines. The cells were cultured in FGM-2 medium supplemented with 5% fetal bovine serum (FBS) in an environment of 5% CO2 at 37°C. The cells were passaged upon reaching confluency and were used at passages 2 to 10. All cell lines used in this study were thoroughly tested to confirm the absence of HIV-1, HBV, HCV, Mycoplasma, bacteria, yeast, and fungal contaminants.

### shRNA and lentivirus production

Epac1 (RAPGEF3) shRNA (TRCN0000047228) cloned into the pLKO.1 lentiviral expression vector, was obtained from Horizon Discovery. Lentiviral particles encapsulating puromycin-resistant lentiviral plasmids were produced using a third-generation system. The constructs and viral packaging plasmids (pVSVG, GAG-POL, TAT, and REV) were co-transfected into HEK 293T cells at 75% confluence using Lipofectamine 2000 (Invitrogen Life Technologies), according to the manufacturer’s instructions. Following transfection, the medium was collected and filtered at 72 h post-infection. NHL and IPF-FBs were infected with lentiviral particles for 72 h and selected with 10 μg/mL puromycin for at least 96 h to ensure stable integration and effective selection.

### Cell Proliferation

Fibroblast proliferation was assessed using a 5-bromo-2’-deoxyuridine (BrdU) incorporation assay (colorimetric) from Roche (Indianapolis, IN, USA) for 48 h following the manufacturer’s instructions.

### Total RNA Isolation, cDNA preparation, and Quantitative RT‒PCR analysis

Total RNA was isolated from the upper lobe of the right lung tissue. The TRIzol reagent (Invitrogen) was used for RNA extraction. Subsequently, cDNA was synthesized using a cDNA Synthesis Kit (Applied Biosystems, Foster City, CA, USA) according to the manufacturer’s instructions. Quantitative RT-PCR was conducted using the PerfeCTa SYBR^TM^ Green FastMix Kit (Quantabio) following the manufacturer’s protocol. The relative comparison method was used to determine fold changes in gene expression. Gene expression was normalized to that of GAPDH, which served as the internal endogenous control. Primer sequences used are listed in **Supplementary Table 4.**

### Immunoblot analysis

To analyze the protein content, cell lysates were prepared using RIPA lysis buffer (Invitrogen) supplemented with a protease inhibitor cocktail (Roche) and a phosphatase inhibitor cocktail (Sigma‒Aldrich). After 20-minute centrifugation at 15000 × g, protein concentrations were determined using the bicinchoninic acid (BCA) assay (Sigma‒Aldrich). The proteins were then separated using SDS‒ polyacrylamide gel electrophoresis (PAGE) and transferred onto polyvinylidene difluoride membranes. The membranes were blocked with 5% skim milk or 3% bovine serum albumin according to the respective antibody specifications. The membranes were then incubated overnight at 4°C with the primary antibodies listed in **Supplementary Table 3**. Next, the membranes were incubated with the corresponding HRP-conjugated secondary antibodies (Cell Signaling Technology). Finally, the luminescence signal was visualized using an enhanced chemiluminescence (ECL) kit (Thermo Fisher Scientific).

### Epac1-based Intramolecular Bioluminescence Resonance Energy Transfer (BRET) assay

The pcDNA3-EBS eukaryotic expression vector was derived from the pQE30-CAMYEL prokaryotic expression vector. The Epac1-BRET sensor (EBS) CAMYEL utilizes a circularly permuted version of Citrine (Citrine cp229, λmax excitation = 480 nm /λmax emission = 535 nm), an enhanced variant of YFP, and Renilla luciferase (RLuc, λmax excitation = 430 nm /λmax emission = 480 nm) as the BRET pair, with human Epac1 (amino acids 149–881) inserted in between, as previously described. When Epac1 is inactive, there is a transfer of energy between the complex luminescent excited enzyme/substrate donor (RLuc/Coelenterazine) and fluorescent molecular acceptor (Citrine). When activated, energy transfer decreases due to a change in the conformation of Epac1, which increases the distance between the two fluorochromes, thus increasing the BRET ratio of RLuc/Citrine. IPF-FBs were transfected with the CAMYEL-EBS construct using Lipofectamine 2000 according to the manufacturer’s instructions. Cells were lysed 48h post-transfection in lysis buffer (HEPES 40 mM, KCl 140 mM, NaCl 10 mM, MgCl2 1,5 mM, Triton 0,5%). Free cell lysates were treated with 2 µM Coelenterazine and AM-001 (20 µM) for ten minutes. The fluorescence from RLuc and yellow fluorescence of the acceptor Citrine-cp229 were measured using a TECAN infinite F500 for 5 min. The cells were then treated with cAMP (100 µM) for 10 min, and fluorescence was measured again over 10 min. The BRET ratio was calculated as RLuc (480nm) / Citrine (535nm), and the basal fluorescence was subtracted after cAMP exposure.

### Neddylation Assay

The neddylation assay was performed in IPF-FBs by transfecting HA-NEDD8 plasmid (2 μg) into the cells. After 24 h of transfection, the cells were treated under the indicated conditions, including TGF-β alone (5 ng/mL, 48 h) or in combination with MG132 (10 μM, 6 h). The cells were lysed using a buffer containing 25 mM Tris-HCl pH 7.4, 150 mM NaCl, 1% NP-40, 1 mM EDTA, 5% glycerol, and protease and phosphatase cocktail inhibitors (Roche). Protein lysates (1 mg) were incubated with the appropriate IgG control antibody (e.g., 0.25 μg/mL) and protein A/G agarose beads at 4°C for 1 h. The samples were centrifuged at 1,000 x g for 1 min at 4°C, and the supernatants were transferred to a clean microcentrifuge tube. The samples were then incubated with a primary HA-Tag antibody (Cell Signaling, 1:50) overnight at 4°C with rotation. Appropriate agarose beads were then added and incubated at 4°C with rotation for 2 h. The samples were centrifuged at 1,000g for 1 min at 4°C, and the supernatant was discarded. Beads were washed with ice-cold RIPA buffer (or phosphate-buffered saline, PBS) three times for 15 min each with constant rotation at 4°C. The samples were then subjected to immunoblotting as previously described.

### Intratracheal BLM mouse model

Animal experiments and handling procedures were performed in accordance with the National Institutes of Health Guide for the Care and Use of Laboratory Animals. Approval was obtained from the Institutional Animal Care and Use Committee of the Icahn School of Medicine at Mount Sinai. The mice were anesthetized by intraperitoneal injection of xylazine and ketamine, carefully immobilized in the supine position on a tray, and intubated using a 20 G angiocath. The tray was then inclined at a 45-degree angle, and the IA-1C microsprayer tip (PennCentury, Wyndmoor, PA, USA) was inserted through the lumen of the angiocath. Subsequently, 50 µL of bleomycin was administered at a dose of 4 units/kg. Once delivery was complete, the microsprayer tip was withdrawn, and the animals were extubated and carefully returned to their respective cages.

### Heart Hemodynamic Studies

Mice were anesthetized with 2-4% isoflurane, intubated via tracheotomy, and mechanically ventilated with 1-2% isoflurane and oxygen (tidal volume: 6 mL/kg; respiratory rate: 100 breaths per minute). The thoracic cavity was opened via a sternotomy to access the organs. After opening the pericardium and fully exposing the heart, an ultrasonic flow probe (2.5 S176; Transonic Systems Inc., Ithaca, NY) was inserted into the right ventricle (RV) to measure right ventricular systolic pressure (RVSP). Hemodynamic data were recorded using a Transonic Scisense PV Control Unit.

### Right ventricular weight measurements

After collecting hemodynamic data, the mice were euthanized to harvest the heart and lungs. The heart was removed from the chest cavity and perfused with PBS to remove blood and clots. The atria and connecting vessels were dissected. The RV was then separated from the heart and weighed. Subsequently, the left ventricle (LV) and septum were weighed. The Fulton Index, illustrating RV hypertrophy, was calculated as the weight ratio of the RV to the LV plus septum (RV weight / [LV + septum weight]).

### Hematoxylin and eosin and Masson’s trichrome staining

RV and Lung tissues were collected, infused with a PBS/OCT mixture (50:50), and subsequently fixed by freezing in OCT at −80°C. Sections (8 µm) were carefully sliced and affixed to ColorFrost glass slides (ThermoFisher Scientific). For histological analysis, lung tissue sections were subjected to hematoxylin and eosin (H&E) and Masson’s trichrome staining (Sigma‒Aldrich). Images were obtained using a light microscope. ImageJ software was used to measure medial thickness and collagen deposition. The Ashcroft score was used to assess pulmonary fibrosis using a validated semi-quantitative approach. This scoring system ranges from 0 (normal lung) to 8 (complete fibrous obliteration of the field). Average scores were calculated using five different sections.

### Immunostaining

Human and mouse paraffin lung sections (8 μm) were deparaffinized and incubated with a blocking solution containing 10% normal goat serum for 1 h at room temperature. The sections were incubated overnight at 4°C with specific antibodies against Epac1 (1:100), FoxO3a (1:50), NEDD8 (1:100), and alpha-SMA (1:250). The primary antibodies used are listed in **Supplementary Table 3**. After washing thrice with PBS, the sections were incubated with secondary antibodies coupled to Alexa Fluor® 488 or Alexa Fluor® 568 (1:500, Invitrogen) for 1 h at 37°C. The cover slides were mounted using a Vectashield mounting medium with DAPI (Vector Laboratories).

### Wheat Germ Agglutinin (WGA) Immunostaining

RV sections were fixed and incubated in cold acetone for 20 min. Following fixation, sections were incubated with a blocking solution containing 10% normal goat serum for 1 h at room temperature. The RV sections were stained overnight at 4°C with fluorescence-tagged wheat germ agglutinin (WGA) (Invitrogen). Imaging was performed using a Zeiss Observer Z.1 microscope (Carl Zeiss) at 160x magnification. The outlines of the cardiac myocytes were traced, and the cardiomyocyte area was analyzed using ImageJ software.

### mRNA sequencing and transcriptomic data analysis

For mRNA sequencing, RNA was extracted from NHL-FBs and IPF-FBs using the QIAGEN RNeasy Mini Kit (Qiagen) following the manufacturer’s protocol. The samples were assessed using a NanoDrop spectrophotometer and Agilent 2100 Bioanalyzer. The integrity and purity of the samples were confirmed by agarose gel electrophoresis. Only samples meeting the criteria of an RNA integrity number (RIN) above 7, OD260/280≥2, and OD260/230 ≥ 2 were used for RNA-seq. Illumina sequencing was conducted using Novogene (Sacramento, CA, USA) on an Illumina NovaSeq 6000 platform. The sequencing strategy involved 250–300 bp insert cDNA libraries with a paired-end 150 bp sequencing approach. For biological triplicate samples, differential expression analysis between the two conditions/groups was performed using the DESeq2 R package. P-values were adjusted using the Benjamini-Hochberg method to control for false discovery rates.

### Bioinformatics and Data Visualization

Data analysis and visualization were performed using ClusterGrammer (accessible at http://amp.pharm.mssm.edu/clustergrammer/). ClusterGrammer is a web-based tool that facilitates interactive exploration and analysis of high-dimensional data through hierarchically clustered heatmaps. Volcano plots were generated to outline the distribution of differentially expressed genes. The x-axis displays log2-fold changes, and the y-axis shows the log10-transformed corrected P-values. The significance threshold was set at 0.05, and an absolute log2-fold change threshold of 1.5 indicated differentially expressed genes. Before heatmap visualization, the raw gene counts were normalized using the logCPM method. The top 2500 genes with the most variable expression were chosen after filtering, and Z-score transformation was applied.

### Immune phenotyping by Flow Cytometry

After systemic perfusion with 10 mL of PBS through the right ventricle, mice lungs were collected, cut into small pieces (<0.5 cm), and enzymatically digested in 10 mL of RPMI (Gibco, Waltham, MA) containing 5% FBS (ATCC, Manassas, VA), 0.5 mg/mL DNase I (grade II, from bovine pancreas; Roche, Penzberg, Germany), and 0.2 mg/mL collagenase IV (Gibco, Waltham, MA) for 1h at 37°C. The samples were passed through a 70 µm cell strainer, washed twice with RPMI-1640 (Thermo Fisher Scientific, Waltham, MA), and further purified by density gradient using a Percoll gradient (Pharmacia, Piscataway, NJ, USA). Single-cell suspensions were labeled for viability (Live/Dead blue; Thermo Fisher Scientific), washed in FACS buffer (2% FBS and 2 mM EDTA in PBS), and Fc receptors were blocked with anti-CD16/CD32 (clone 93; BioLegend, San Diego, CA, USA). Staining for surface markers was performed on ice for 45 min, washed with FACS buffer (2% FBS and 2 mM EDTA in PBS), and fixed/permeabilized using the FOXP3/Transcription Factor Staining Buffer Set (Thermo Fisher Scientific). Finally, the samples were labeled with intracellular and extracellular markers on ice for 45 min. Samples were acquired using a 5-laser Cytek Aurora cytometer (Cytek Biosciences, Fremont, CA) and analyzed using FlowJo 10.8.2 software (BD Biosciences, Franklin Lakes, NJ). The antibodies used are provided in **Supplementary Table 5**.

### Statistical analysis

Each experiment was conducted in triplicate to ensure robustness. The results are expressed as mean ± standard error of the mean (SEM). The unpaired t-test was used to compare the two conditions. In cases where comparisons involved more than two conditions, a one-way analysis of variance (ANOVA) was conducted, followed by Tukey’s correction for post hoc comparisons. All statistical analyses were performed using the GraphPad Prism software (GraphPad Software, Inc., La Jolla, CA, USA).

## RESULTS

### Epac1 expression is upregulated in the lungs of patients with IPF and mice challenged with BLM

To determine the contribution of Epac proteins to PF pathogenesis, we first examined Epac1 expression in the lungs of patients with IPF (65 ± 10 years old, *n*=8) and matched healthy control donors (62 ±10 years old, *n*=8). We found that Epac1 mRNA expression level was higher in the IPF lungs than in healthy controls (**Figure 1A**), whereas Epac2 expression did not change (**Supplementary Figure 1A**). Additionally, co-immunostaining showed that SMA^+^ fibroblasts (green) strongly expressed Epac1 (red) in fibrotic areas and fibroblastic foci in IPF lung samples compared with healthy controls (**Figure 1B**). Similarly, increased Epac1 protein levels were observed in proliferating isolated IPF fibroblasts (IPF-FBs) expressing α-SMA and Cyclin D1 compared to those in normal human lung fibroblasts (NHL-FBs) (**Figure 1C & Supplementary Figure 1B**). Finally, Epac1 expression was also increased following a single intratracheal instillation of BLM (4 U/kg) in mice after 28 days^22^ (**Supplementary Figure 1C, D**). These data suggest that increased Epac1 expression is a feature of PF and led us to investigate the role of Epac1 in PF development.

**Figure 1.**
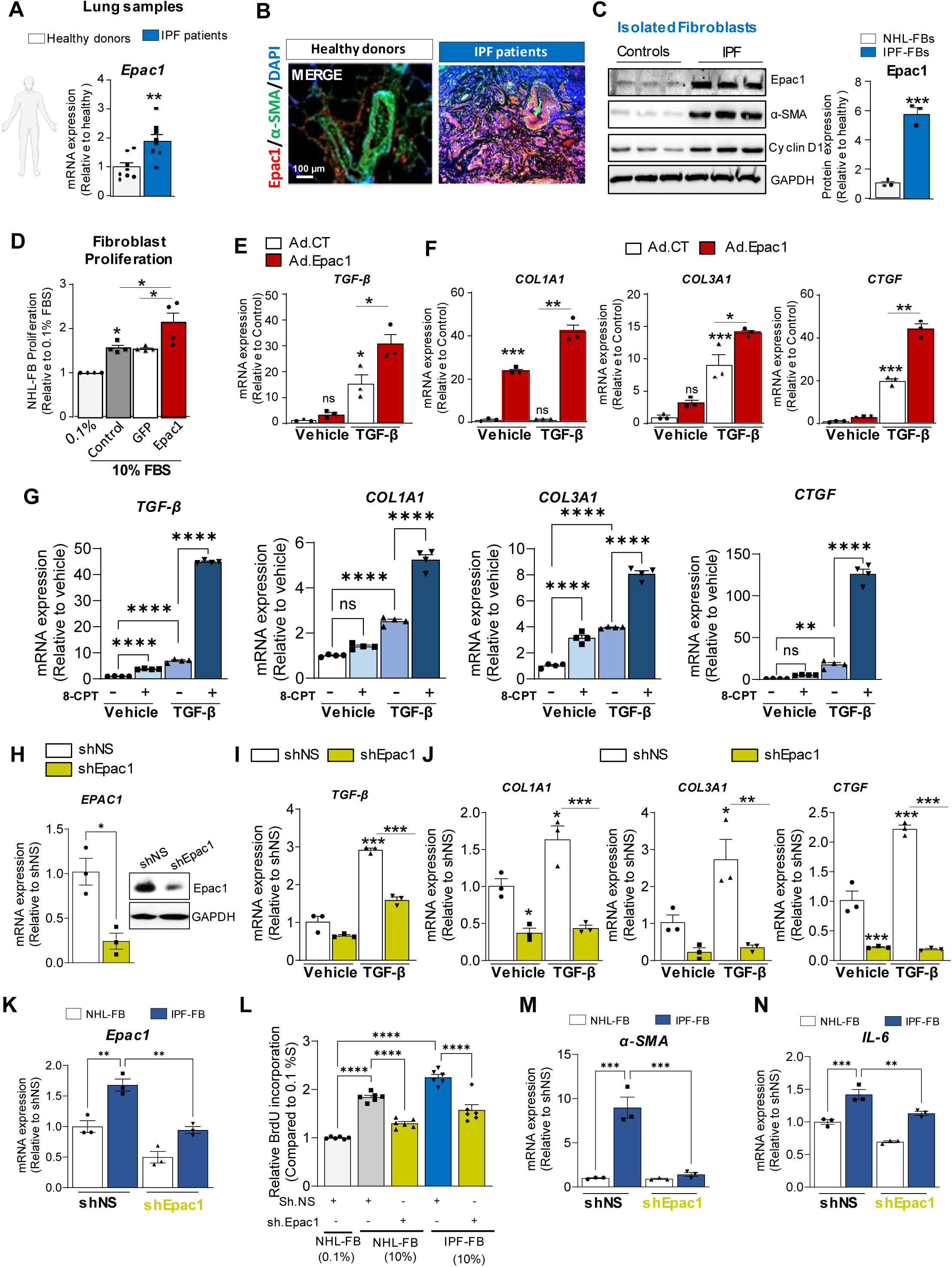
Epac1 Expression in IPF and fibrosis-related genes and proliferation in lung fibroblasts. **A.** Epac1 (left panel) mRNA levels were assessed in lung tissues from IPF patients (*n* = 8) and healthy control lungs (*n* = 8) using qRT-PCR. **B.** Representative immunofluorescence staining for Epac1 and α-smooth muscle actin (α-SMA) in lung tissues from healthy donors and IPF patients. Scale bar-100 μm. **C.** Representative immunoblot (left panel) and quantification (right panel) of Epac1 normalized using GAPDH in isolated IPF-FB (*n* = 3) and normal human lung fibroblasts, NHL-FBs (*n* = 3). **D**. Proliferation was measured in media supplemented with low serum (0.1%) or high serum (10% FBS) concentrations using a BrdU assay for 72 hours in NHL-FBs overexpressing adenovirus (Ad).Epac1 and Ad.GFP (Ad.CT). **E**. TGF-β mRNA expression in NHL-FBs infected with Ad.CT or Ad.Epac1 alone or co-treated with TGF-β (5 ng/mL) for 48 hours. **F**. Profibrotic markers COL1A1, COL3A1, and CTGF mRNA expression levels in NHL-FBs overexpressing either Ad.CT or Ad.Epac1 with or without TGF-β treatment for 48 hours. **G**. mRNA expression levels of TGF-β and profibrotic markers in NHL-FBs co-treated with 8-CPT (5 ng/mL) and TGF-β (5 ng/mL). **H**. Epac1 mRNA and protein expression were measured by qRT-PCR and western blot, respectively, in NHL-FBs overexpressing either a non-silencing shRNA (shNS) or a specific shRNA against Epac1 (shEpac1). **I-J**. Analysis of TGF-β, COL1A1, COL3A1, and CTGF mRNA expression in NHL-FBs overexpressing either shNS or sh.Epac1 treated with vehicle or TGF-β for 24 hours. **K**. Epac1 mRNA expression in NHL-FBs or IPF-FBs overexpressing shNS or sh.Epac1. **L**. NHL-FBs and IPF-FB proliferation were assessed using a BrdU assay under the specified experimental conditions. **M-N**. α-SMA and IL-6 mRNA expression levels in NHL-FBs and IPF-FBs overexpressing shNS or shEpac1. The data are presented as the mean ± SEM; *p < 0.05, **p < 0.01, ***p < 0.001.

### Epac1 regulates primary normal human lung fibroblasts and IPF fibroblasts growth and activation

Lung FBs are central mediators of pathological fibrotic accumulation in the ECM and their proliferation occurs in response to prolonged injury and inflammation^8^. We first used a gain-of-function approach with an adenovirus encoding Epac1 (Ad.Epac1) to assess the effects of Epac1 overexpression on activated human normal lung FB (NHL-FB) proliferation and profibrotic gene expression. As expected, Ad.Epac1-induced Epac1 overexpression with no changes in Epac2 expression (**Supplementary Figure 2A, B**). Interestingly, Ad.Epac1 potentiated NHL-FB proliferation cultured in high serum concentrations (10%FBS) (**Figure 1D**). Moreover, Epac1 overexpression potentiated TGF-β mRNA levels in TGF-β1-stimulated NHL-FBs (**Figure 1E**), and the expression of profibrotic genes, such as collagen type 1 alpha 1 (COL1A1), collagen type 3 alpha 1 (COL3A1), and connective tissue growth factor (CTGF) (**Figure 1F**). These data suggest that Epac1 is a key regulator of lung FB growth and pathway activation under stress stimuli.

Next, NHL-FB cells were stimulated with the Epac1-specific agonist, 8-pCPT-2′-O-Me-cAMP (8-CPT), to confirm these results. NHL-FB stimulation with 8-CPT promoted the expression of TGF-β and profibrotic genes, such as COL1A1, COL3A1, and CTGF (**Figure 1G**). Markedly, the addition of TGF-β1 significantly potentiated the effects of 8-CPT on fibrosis-related gene expression (**Figure 1G**). The potent stimulatory effect of TGF-β1 in the presence of 8-CPT suggests that the secretion of TGF-β1 from the surrounding cells stimulates lung FB and may lead to increased Epac1 expression and/or activity to induce profibrotic markers. Finally, we validated the robustness of these results using shRNA-mediated Epac1 knock-down (sh.Epac1) in NHLF-FBs without change in Epac2 expression **(Figure 1H & Supplementary Figure 2C**, respectively). Surprisingly, sh.Epac1 reduced TGF-β1 mRNA **(Figure 1I)**, and COL1A1, COL3A1 and CTGF levels **(Figure 1J**) in NHLF-FBs stimulated with TGF-β. Remarkably, IPF-FBs expressed Epac1 at higher mRNA levels compared to NHL-FBs (**Figure 1K**), and Epac1 knock-down diminished the anti-proliferative activity of NHL-FBs and IPF-FB cultured in 10% FBS (**Figure 1L**). Notably, α-SMA and IL-6 mRNA levels were reduced in Epac1-depleted IPF and NHL-FBs (**Figure 1M, N)**. Altogether, these data demonstrate that Epac1 regulates the growth and activation of primary human lung FBs.

### Selective pharmacological inhibition of Epac1 with AM-001 prevents lung fibroblast proliferation and the expression of profibrotic markers *in vitro*

The inhibition of Epac1 using AM-001 **(Supplementary Figure 3A)** effectively blocked cAMP-induced Epac1 activation in IPF-FB as assessed using an Epac1-based BRET sensor (**Figure 2A**). Notably, AM-001 significantly reversed the stimulating effect of high serum and TGF-β1 on NHL-FB proliferation (**Figure 2B**) as well as the effects of TGF-β-induced increases in TGF-β1, COL1A1, COL3A1, and CTGF mRNA expression (**Figure 2C**). As expected, we found that CE3F4^37^, another pharmacological inhibitor of Epac1, mimicked the effect of AM-001 on NHL-FB proliferation and induction of fibrosis markers (**Supplementary Figure 3B-D)**. These results demonstrate that selective Epac1 pharmacological inhibition prevents the activation of lung FB growth.

**Figure 2.**
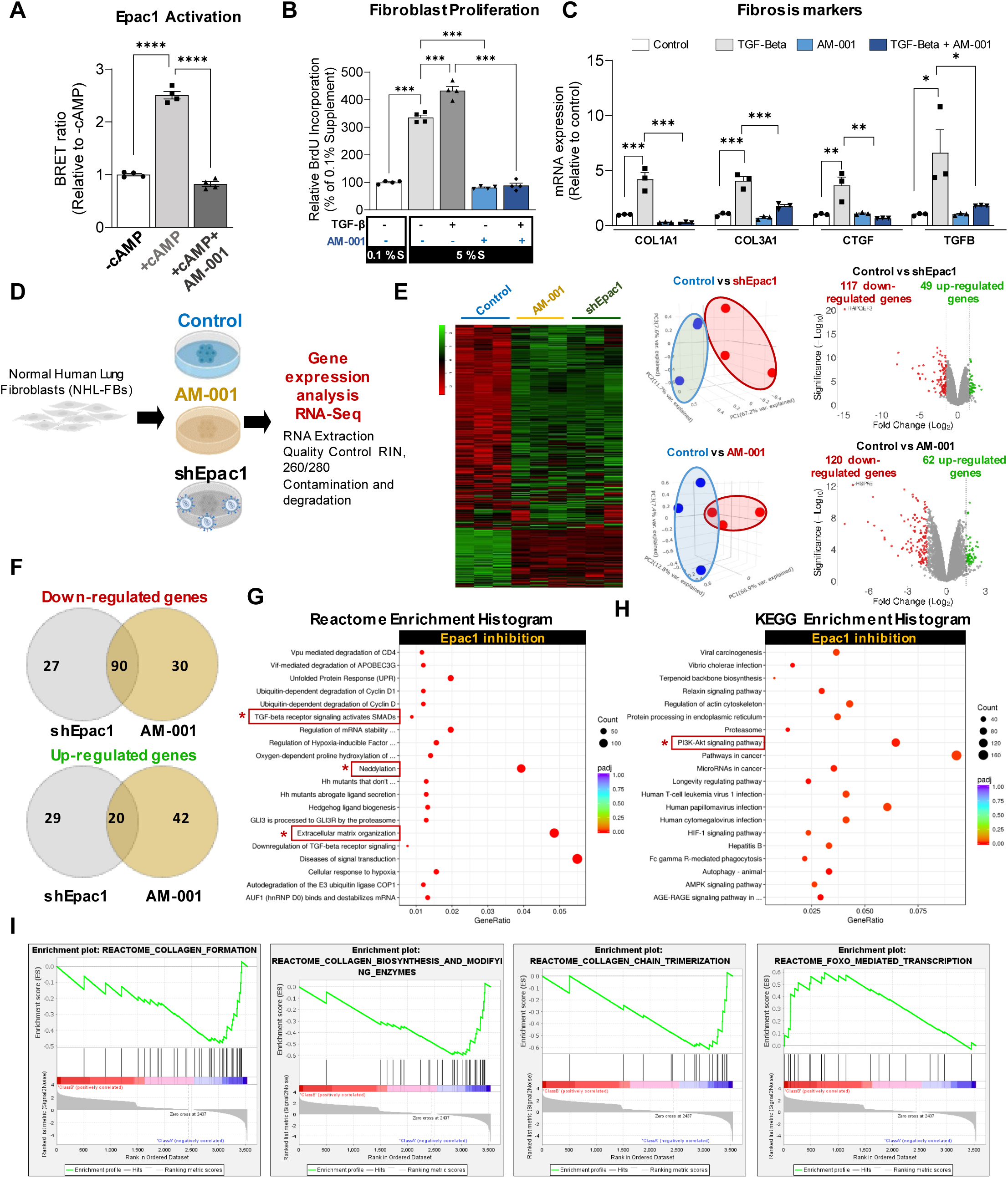
Pharmacological inhibition of Epac1 by AM-001. **A.** Epac1 activity using the BRET-CAMYEL sensor in IPF-FBs stimulated with cAMP alone or in combination with AM-001. **B.** NHL-FBs proliferation treated with TGF-β alone or in combination with AM-001 in media supplemented with low serum (0.1%) or high serum concentrations (10%), and cell proliferation was measured using a BrdU assay. **C.** Expression of fibrosis markers mRNA levels of TGF-β, COL1A1, COL3A1, CTGF in NHL-FBs pre-treated with AM-001 or DMSO (as vehicle control) for 24 hours, followed by activation by TGF-β for an additional 24 hours. Quantification was performed using qRT-PCR. **D.** Experimental design for RNA sequencing (RNA-seq) of NHL-FBs treated with AM-001 for 48 hours or infected with a specific shRNA against Epac1 (sh.Epac1) or sh.NS for 72 hours. **E.** Heatmap and volcano plot displaying genes that were either upregulated or downregulated in response to AM-001 or sh.Epac1 compared to that in control cells. **F.** Venn diagrams showing the overlap of genes commonly downregulated and upregulated by AM-001 and sh.Epac1 treatment. The upper panel represents the downregulated genes, while the right panel shows the upregulated genes. The intersecting areas highlight the genes affected by both treatments. **G-H**. Reactome and KEGG, pathway enrichment analysis, for genes significantly differentially expressed in response to AM-001 or shEpac1. **I.** An enrichment plot was generated through gene set enrichment analysis (GSEA), focusing on Reactome pathways related to collagen formation, collagen biosynthesis and modifying enzymes, collagen chain trimerization, and FOXO-mediated transcription. The data are presented as the mean ± SEM; *p < 0.05, **p < 0.01.

### Characterization of Epac1 transcriptome involved in profibrotic pathways in NHL-FBs

To globally identify Epac1 target genes, we performed bulk RNA-Seq in proliferating stimulated NHL-FBs treated with AM-001 or shEpac1 for 48 h (**Figure 2D**). A total of 182 and 166 differentially expressed genes (DEG) were identified between the control (CTL)- and activated NHL-FBs and treated with AM-001 or sh.Epac1, respectively (**Figure 2E**). More specifically, the expression of 62 genes was significantly upregulated, whereas that of 120 was downregulated in AM-001 NHL-FB-treated cells (**Figure 2E**). Comparison analysis of the DEGs revealed specific transcriptomic signatures in NHL-FB treated with AM-001 or shEpac1 (**Figure 2F**). As anticipated from the extensive dataset analysis, the results of the Gene Ontology (GO) and REACTOME pathway analysis indicated that critical gene signatures predominantly relate to processes such as ECM and extracellular structure organization. These processes include collagen synthesis, ECM organization, and involvement in fibroblast growth factor receptor 1 (FGFR1), tenascin C (TNC), TGF-β signaling, and neddylation pathway (**Figure 2G**). Furthermore, Kyoto Encyclopedia of Genes and Genomes (KEGG) pathway analysis showed that the downregulated DEGs were strongly associated with pathways, including the proteasome and PI3K-AKT signalosome (**Figure 2H**). As predicted by Gene Set Enrichment Analysis (GSEA), most genes with downregulated expression following Epac1 inhibition were associated with collagen formation, biosynthesis, and trimerization (**Figure 2I**). Interestingly, we identified Forkhead box O 3 (FoxO3a), as the top transcription factor (**Figure 2I**) that has been shown to reduce lung fibrosis by inhibiting FB activation and reducing ECM deposition ^38^.

### Epac1 inhibition with AM-001 prevents TGF-β1-induced profibrotic STAT3, SMAD2/3, and AKT signaling in NHL-FBs and IPF-FBs

Treatment of IPF-FBs with the TGF-β1 receptor inhibitor SB431542 (10μM) significantly reduced Epac1 mRNA levels compared to the nonactivated or non-TGF-β1-stimulated NHL-FBs (**Figure 3A**), confirming that TGF-β1 induced Epac1 expression in IPF-FBs. In addition, AM-001 exerted a strong anti-proliferative effect on serum-induced IPF-FB proliferation (**Figure 3B**).

**Figure 3.**
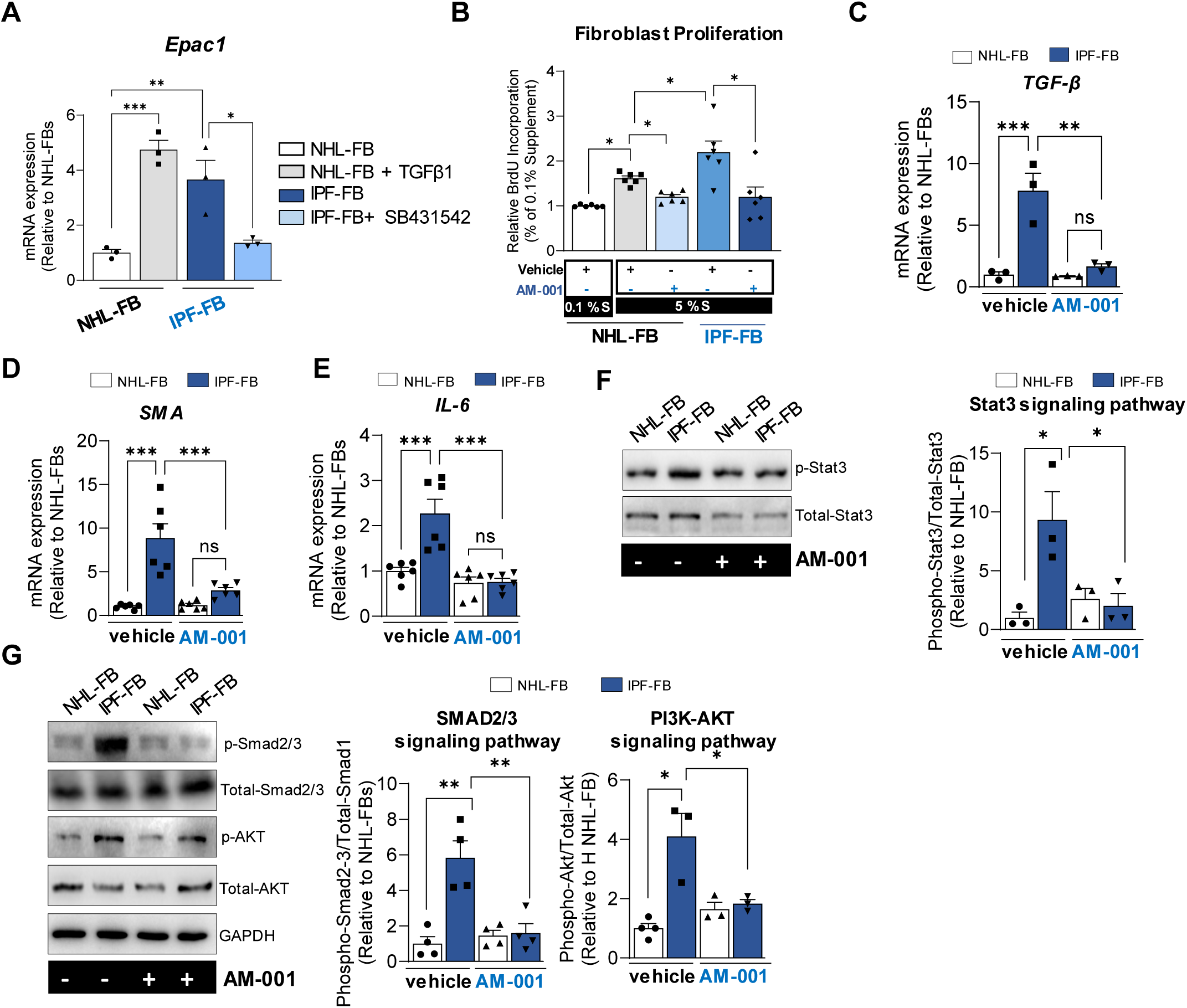
Effects of pharmacological inhibition of Epac1 in IPF-FBs on the STAT3, PI3K/AKT, and SMAD2/3 pathways. **A.** Epac1 mRNA levels in normal human lung fibroblasts (NHL-FBs) and fibroblasts isolated from idiopathic pulmonary fibrosis patients (IPF-FBs). NHL-FB or IPF-FBs were treated for 48 hours with either TGF-β or SB431542 (10 µM), a potent and selective inhibitor of the TGF-β type I receptor. **B.** Proliferation was assessed by BrdU labeling in both NHL-FBs and IPF-FBs after treatment with AM-001 in the presence of low- or high-serum medium concentrations (10%) for 72 hours. **C-E.** mRNA expression of the levels of TGF-β (**C**), α-SMA (**D**), and IL-6 (**E**) in nonactivated NHL-FBs and IPF-FBs treated with either vehicle or AM-001 for 48 hours. Quantification was performed using qRT-PCR. **F.** Western blot analysis of phospho-STAT3 and total-STAT3 levels in both NHL-FBs and IPF-FBs in the presence or absence of AM-001. The left panel shows a representative immunoblot and the right panel shows the phospho-STAT3/total-STAT3 ratio quantification. **G.** Immunoblot analysis of phospho-SMAD2/3, total SMAD2/3, phospho-AKT, total AKT, and GAPDH (as loading controls) in NHL-FBs and IPF-FBs treated with vehicle or AM-001. The left panel shows representative immunoblots and the right panel shows the quantification of the SMAD signaling pathway (phospho-Smad2-3/total-Smad1) and the PI3K-AKT signaling pathway (phospho-AKT/total-AKT).

Next, we evaluated the effects of AM-001 on Epac1 and TGF-β1 transcripts in IPF-FBs. We first found that Epac1 mRNA levels were decreased only in AM-001-treated IPF-FBs compared to nonactivated non-TGF-β1-stimulated NHL-FBs (**Supplementary Figure 5A**). Moreover, TGF-β1 mRNA levels were higher in IPF-FBs than in NHL-FBs, and AM-001 treatment abolished this effect without affecting basal TGF-β1 levels in non-stimulated NHL-FBs (**Figure 3C**). Taken together, these results suggested that reduced TGF-β levels may restore Epac1 expression. Interestingly, AM-001-treated IPF-FBs significantly attenuated the mRNA expression of α-SMA and IL-6 (**Figure 3D, E**). Consequently, the protein levels of p-STAT3, p-SMAD2/3, and p-AKT were markedly decreased in IPF-FBs treated with AM-001 (**Figure 3F, G**). Our results showed that Epac1 simultaneously activates pro-inflammatory and profibrotic signaling pathways related to IL-6/STAT3, TGF-β/SMAD2/3, and AKT signaling in IPF-FBs. Next, we analyzed our RNA-Seq dataset to identify the top 50 transcription factors using the Encyclopedia of DNA Elements (ENCODE) and ChIP Enrichment Analysis (ChEA) libraries and identified the nuclear factor FoxO3a as a potential effector of Epac1 profibrotic signaling pathways. Previous studies have shown that FoxO3a is under the control of the PI3K/AKT signaling pathway and is implicated in the differentiation of lung FBs and the hyperproliferation phenotype ^34, 39^. Consistent with these studies, we found that Epac1 overexpression reduced FoxO3a mRNA level in NHL-FBs **(Supplementary Figure 4A),** whereas Epac1 silencing reversed this effect in IPF-FBs without effecting FoxO3a expression in non-activated NHL-FBs (**Supplementary Figure 4B**), suggesting that Epac1 regulates FoxO3a expression level.

### AM-001 blocks endogenous FoxO3a degradation through neddylation in IPF-FBs

Neddylation-like ubiquitination is a reversible post-translational modification that plays a crucial role in controlling substrate degradation and is catalyzed by the successive enzymatic cascade of NEDD8-activating enzyme E1 (NAE1), NEDD8-conjugating enzyme E2, and substrate-specific NEDD8-E3 ligases^40–42^. Based on our RNAseq analysis that identified neddylation-related genes and FoxO3a as being up- and downregulated in NHL-FBs, respectively (**Figures 2G, I**), we next examined the effects of AM-001 on the neddylation pathway and FoxO3a levels in AM-001-treated IPF-FBs. Among the neddylation-related genes, NAE1, NEDD8, and UBE2M mRNA levels in IPF-FBs were significantly decreased by AM-001 (**Figure 4A**), as well as UBA3 and UBE2F transcripts (**Supplementary Figure 5B, C**). On the contrary, the expression level of FoxO3a mRNA was restored in AM-001 treated IPF-FBs (**Figure 4B**). Similar reductions in the levels of neddylation-related genes were observed in Epac1-depleted IPF-FBs, although the difference for UBE2F was not significant (**Supplementary Figure 5C-G**). Consistently, we found that TGF-β1 treatment increased both FoxO3a phosphorylation at Thr32 (pFoxO3a^Thr32^) and NEDD8 protein levels in NHL-FBs (**Figure 4C**). In IPF-FBs, the expression levels of pFoxO3a^Thr32^ and NEDD8 were high under basal conditions with decreased FoxO3a expression (**Figure 4C**). These data suggest that NEDD8 may promote FoxO3a degradation when it is phosphorylated and that Epac1 inhibition may reverse this effect.

**Figure 4.**
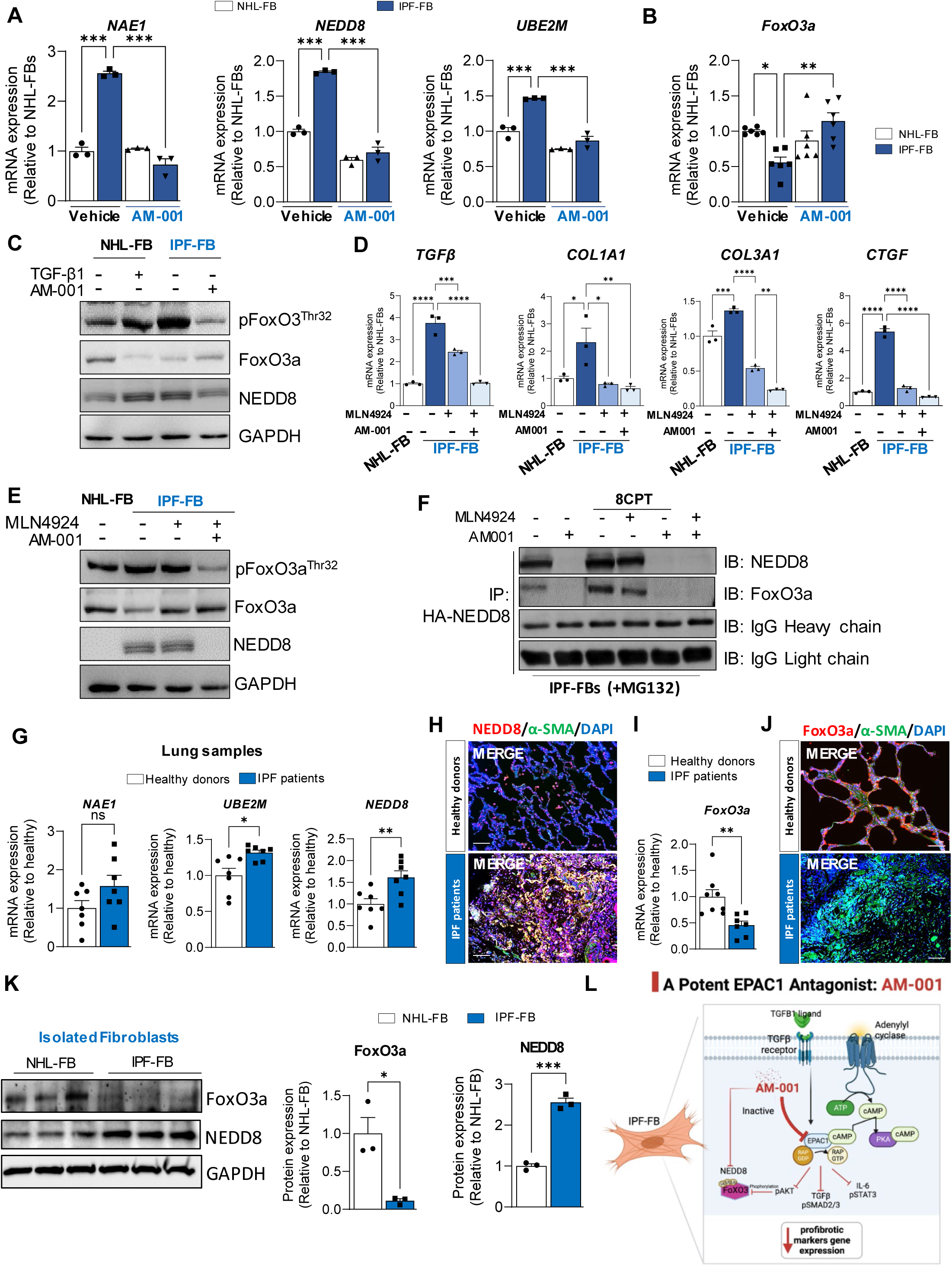
Molecular responses to AM-001 and neddylation inhibition in lung fibroblasts. **A.** Transcript levels of NAE1, NEDD8, and UBE2M in NHL-FBs and IPF-FBs after 48 hours of treatment with either vehicle or AM-001. **B.** FoxO3a mRNA expression in NHL-FBs and IPF-FBs after 48 hours of treatment with either vehicle or AM-001. **C.** Representative immunoblot analysis showing the levels of FoxO3a phosphorylated at Thr32 (phospho-FoxO3a^Thr32^), total FoxO3a, and NEDD8 in indicated conditions. GAPDH was used as a loading control. **D.** The expression mRNA levels of fibrosis markers TGF-β, COL1A1, COL3A1, and CTGF were measured by qRT-PCR in NHL-FBs or IPF-FBs treated with MLN4924 alone, a small molecule inhibitor of NEDD8-activating enzyme (NAE), or in combination with AM-001 for 48 hours. **E.** Representative immunoblot analysis of phospho-FoxO3a^Thr32^, total FoxO3a, and NEDD8 protein levels in NHL-FBs or IPF-FBs treated with vehicles, MLN4924 alone (5 µM) or in combination with AM-001for 48 hours. **F.** Immunoprecipitation of NEDD8 followed by immunoblot analysis of NEDD8 and total FoxO3a in IPF-FBs transfected with HA-NEDD8 and pre-treated with proteasome inhibitor MG132 (10 µM) and under the indicated conditions. **G.** The transcript levels of NAE1, UBE2M, and NEDD8 in lung tissue samples from healthy donors and IPF patients were assessed by RT-qPCR. **H**. Co-immunostaining of NEDD8 (red) and α-SMA (green) in lung tissue sections from both healthy donors and IPF patients. Nuclei were counterstained with DAPI (blue). Scale bar-100 μm. **I-J**. FoxO3a mRNA levels and FoxO3a co-immunostaining (red) and α-SMA (green) in lung tissue sections from both healthy donors and IPF patients. Scale bar-100 μm. **K**. Western blot analysis of FoxO3a and NEDD8 in NHL-FBs and IPF-FBs. The left panel shows a representative immunoblot, and the right panel shows the ratio quantification using GAPDH as loading control. **L.** Pathway schematic of AM-001 effects on Epac1 and the downstream molecular pathway. Data are presented as the mean ± SEM; *p < 0.05, **p < 0.0, ***p < 0.001.

To validate this assumption, we examined whether the ubiquitin-like protein NEDD8 was involved in the increased degradation of FoxO3a proteins in the context of IPF. As expected, NEDD8 protein expression levels were increased in NHL-FBs stimulated with TGF-β and IPF-FBs, whereas AM-001 reversed this effect by decreasing NEDD8 protein levels and restoring total FoxO3a levels in IPF-FBs (**Figure 4C**). In addition, we found that a selective inhibitor of NEDD8-activating enzyme E, MLN4924 (Pevonedistat), mimicked the effect of AM-001 on the expression of profibrotic markers in IPF-FBs (**Figure 4D**). Importantly, co-treatment of IPF-FBs with MLN4924 and AM-001 reduced phospho-FoxO3a^Thr32^ levels and restored the total FoxO3a expression by suppressing NEDD8 protein levels, suggesting that pFoxO3a degradation was regulated by NEDD8-dependent neddylation (**Figure 4E**). Consistently, AM-001 reduced NEDD8-FoxO3a interaction in IPF-FB cells pre-treated with the proteasome inhibitor MG132 (**Figure 4F**). Furthermore, the interaction of endogenous FoxO3a proteins was prevented by the addition of AM-001 (**Figure 4F**), while IPF-FB pretreatment with 8-CPT to activate Epac1 and the addition of MLN4924 partially restored the level of NEDD8 and NEDD8-FoxO3a complex, while AM-001 blocked completely this interaction (**Figure 4F**). These findings suggest that activated Epac1 maintains the NEDD8 expression level, which promotes FoxO3a degradation in IPF-FBs, while AM-001 reverses this effect (**Figure 4F**). Overall, our results suggest that the proteasome-mediated degradation of phospho-FoxO3a is regulated by the NAE1-NEDD8 axis in IPF-FBs.

Accordingly, we demonstrated that the neddylation pathway was upregulated in human IPF. Indeed, UBE2M and NEDD8 mRNA expression but not NAE1 were significantly increased in the lungs of patients with IPF (**Figure 4G, H**), while FoxO3a mRNA level was significantly reduced (**Figure 4I, J**). Of note, we observed abundant NEDD8 protein expression (red) in IPF fibrotic lung tissues co-stained with the myofibroblast marker α-SMA (green), while FoxO3a (red) protein expression was reduced (**Figure 4H, J**). Compared to NHL-FBs, we confirmed that FoxO3a and NEDD8 protein levels were inversely correlated in NHL-FBs and IPF-FBs (**Figure 4K**). Overall, our data suggest that AM-001, by inhibiting Epac1, exerts pleiotropic effects on IPF-FB (**Figure 4L**).

TGF-β regulates the fibroblast growth factor (FGF)/fibroblast growth factor receptor (FGFR) signaling cascade in lung FBs ^43, 44^. It also upregulates the TNC-related pathway^45–47^. FGFR1 and TNC both trigger lung fibrosis^47, 48^ and are upregulated in human IPF lung samples compared with healthy lung samples (**Supplementary Figure 6A, B**). FGFR1 and TNC mRNA expression levels were significantly diminished in AM-001-treated or Epac1-depleted IPF-FBs compared to controls (**Supplementary Figure 7A-D).**

### Epac1 deficiency protects mice from BLM-induced fibrosis

Next, we sought to determine the impact of Epac1 genetic inhibition on PF development after BLM administration. The BLM aerosolization challenge in mice induces lung injury with a subsequent fibroproliferative response, leading to the production of cytokines, such as TGF-β and IL-6^20, 22, 49^. For this, global Epac1 KO mice (14 weeks old) and wild-type (WT) mice were randomly subjected to a single intratracheal (IT) instillation of BLM (4 U/kg) and evaluated 28 days later^22^ (**Figure 5A**). We first validated the significant decrease in Epac1 mRNA and protein expression in the lungs of Epac1 KO mice without significant changes in Epac2 levels (**Figure 5B**). BLM treatment specifically increased lung Epac1 without affecting Epac2 expression in WT mice (**Supplementary Figure 8A, B**). Remarkably, hallmarks of PF formation, such as deposition of ECM, Ashcroft histopathology, TGF-β, and fibrotic markers, were all significantly reduced in BLM-treated Epac1 KO mice compared with BLM-treated WT mice (**Figure 5C-F**).

**Figure 5.**
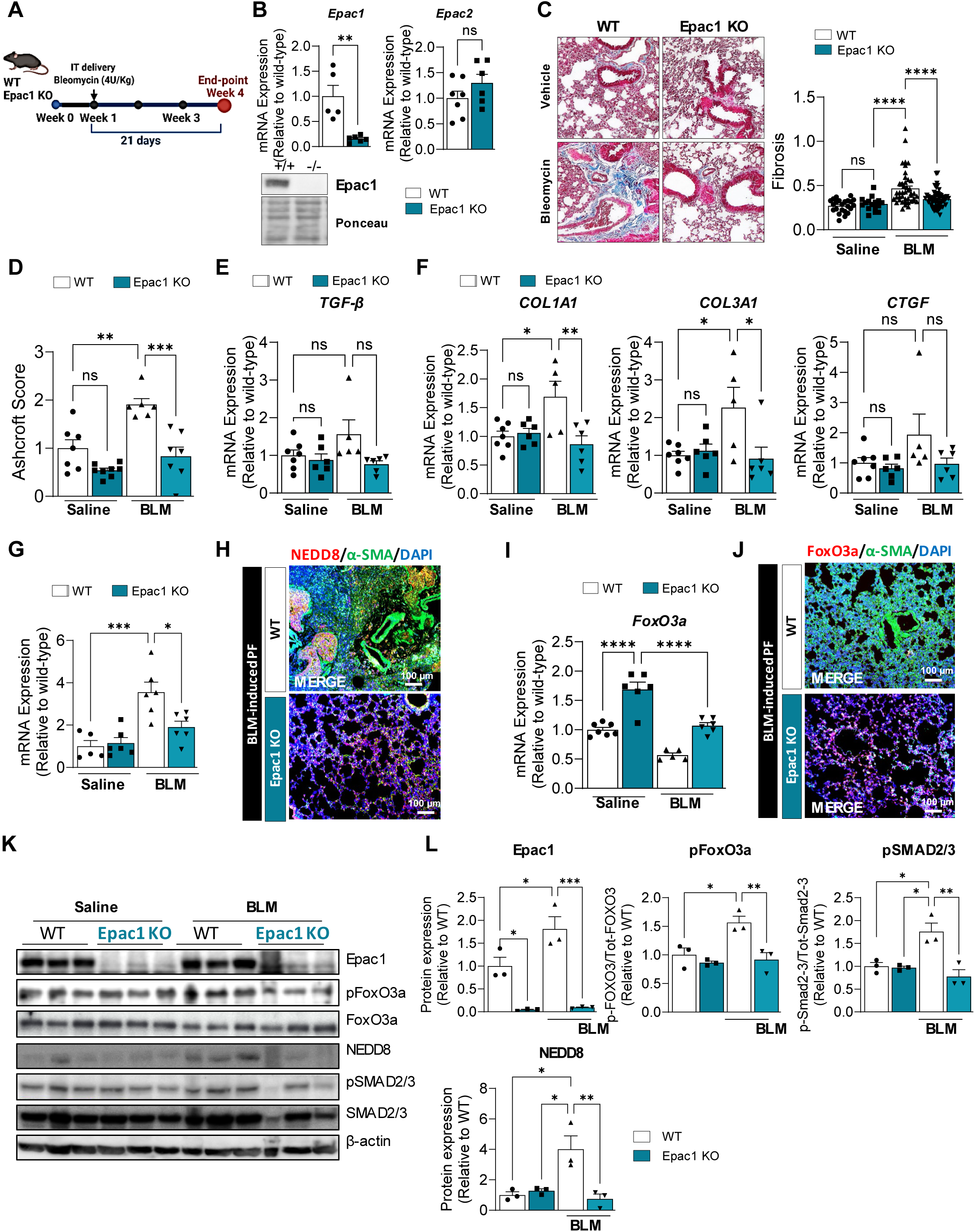
Epac1 deficiency prevents bleomycin-induced lung fibrosis. **A.** Schematic of the experimental design for the induction of bleomycin (BLM)-induced lung fibrosis. Wild-type littermates (WT) and Epac1 knock-out mice (Epac1^-/-^) were subjected to a single intratracheal aerosolized delivery of BLM at a dose of 4 U/kg, while the sham control group received the saline as a vehicle. The animals were sacrificed 28 days after BLM administration. **B.** Epac1 mRNA and protein level, and Epac2 mRNA levels in WT and Epac1^-/-^ mice were assessed by qRT-PCR and western blotting. **C**. Representative images of Masson’s trichrome staining to visualize fibrotic changes of lung sections from WT and Epac1^-/-^ mice challenged with BLM (left panel). The right panel shows the quantification of the percentage of the fibrotic area, expressed as the fold change relative to the saline WT control. **D.** The Ashcroft scoring system was used to assess the severity of lung fibrosis in Epac1^-/-^ relative to that in WT mice challenged with saline. **E-F.** Pro-fibrosis markers TGF-β, COL1A1, COL3A1, and CTGF mRNA expression levels were measured by qRT-PCR in lung tissues from WT and Epac1^-/-^ mice treated with saline (vehicle) or BLM. **G.** NEDD8 mRNA expression in lung tissue was measured by qRT-PCR. **H**. Co-immunostaining for NEDD8 (red) and α-SMA (green) in lung sections from WT and Epac1^-/-^ mice challenged with BLM. Nuclei were counterstained with DAPI (blue). Scale bar-100 μm. **I.** FoxO3a mRNA expression in lung tissue from WT and Epac1^-/-^ mice challenged with vehicle or BLM. **J.** Co-immunostaining for FoxO3a (red) and α-SMA (green) in lung sections from WT and Epac1^-/-^ mice challenged with BLM. Nuclei were counterstained with DAPI (blue). Scale bar-100 μm. **K-L.** Representative immunoblot analysis and quantification of Epac1, NEDD8, phospho-FoxO3a and total FoxO3a, phospho-Smad2/3, and total Smad2/3. β-actin was used as a loading control. The data are presented as the mean ± SEM; *p < 0.05, ***p < 0.001, **p < 0.01.

At the molecular level, FoxO3a levels were higher in BLM-treated Epac1-KO mice than in BLM-treated WT mice (**Figure 5G**). BLM instillation led to a significant decrease in FoxO3a in WT mice, while in Epac1 KO mice, FoxO3a was also decreased, but to a level similar to saline-treated WT control mice (**Figure 5G**). The effect of Epac1 deficiency on FoxO3a expression was further confirmed by immunofluorescence staining (**Figure 5H**). Remarkably, NEDD8 mRNA and protein expression levels assessed by qRT-PCR and immunostaining were significantly decreased compared to BLM-treated WT mice (**Figure 5I, J**). Finally, we quantified NEDD8 and pFoxO3a and p-SMAD2/3 levels by immunoblotting and observed significantly elevated NEDD8 protein levels as well as pFoxO3a^Thr32^ and p-SMAD2/3 in BLM-treated WT mice but lower in BLM-treated Epac1 KO mice (**Figure 5K, L)**. These results suggest that the regulatory effect of Epac1 on NEDD8 and FoxO3a expression observed *in vitro* in IPF-FBs level also occurs *in vivo*. Accordingly, FGFR1 and TNC mRNA expression was downregulated in BLM-treated Epac1 KO mice (**Supplementary Figure 8C, D**). Altogether, these data demonstrate that Epac1 plays a role in the development of lung fibrosis and that its inactivation confers marked protection against PF.

### Therapeutic effect of AM-001 in PF progression

Next, we tested the effects of AM-001 in a murine model of BLM-induced PF. BLM-challenged mice were randomly assigned to receive either vehicle or AM-001 (10 mg/kg) intraperitoneally (IP) every alternate day for 14 days (**Figure 6A**). We found that AM-001 treatment profoundly reduced fibrotic lesions in BLM-treated animals compared with vehicle treatment (**Figure 6B**) and significantly lowered the mRNA levels of profibrotic markers (**Figure 6C**). Remarkably, AM-001 treatment reversed the BLM-induced effects by significantly decreasing the mRNA expression levels of NAE1, UBE2M, and NEDD8 (**Figure 6D-F)**. Consistently, the results of co-immunostaining and immunoblotting revealed that AM-001 lowered NEDD8 (red) and restored FoxO3a (red) (**Figure 6G, H**) while decreasing Epac1, p-FoxO3a, and p-SMAD2/3 in the lungs of BLM-treated animals compared to those in the vehicle-treated group (**Figure 6H, I**). In addition, AM-001 reduced FGFR1 expression and tended to decrease TNC mRNA levels (**Supplementary Figure 9A, B**). Next, we assessed the effect of AM-001 in pulmonary vascular remodeling secondary to BLM-induced pulmonary fibrosis ^50^. Interestingly, compared to vehicle-treated BLM mice, AM-001 reversed distal pulmonary artery vascular remodeling, as assessed by morphometric analysis (**Supplementary Figure 10A, B)**, and significantly alleviated right ventricular (RV) remodeling by decreasing RV systolic pressure (RVSP) and RV hypertrophy, as revealed by a reduced Fulton Index, as well as the downregulation of hypertrophic genes (**Supplementary Figure 11A-C**). Finally, histological analysis showed a reduction in RV fibrosis and cardiomyocyte size when treated with AM-001 compared to the vehicle (**Supplementary Figure 11D-G**). Taken together, these results demonstrate that the administration of the Epac1 antagonist AM-001 protects against PF progression and subsequent RV remodeling.

**Figure 6.**
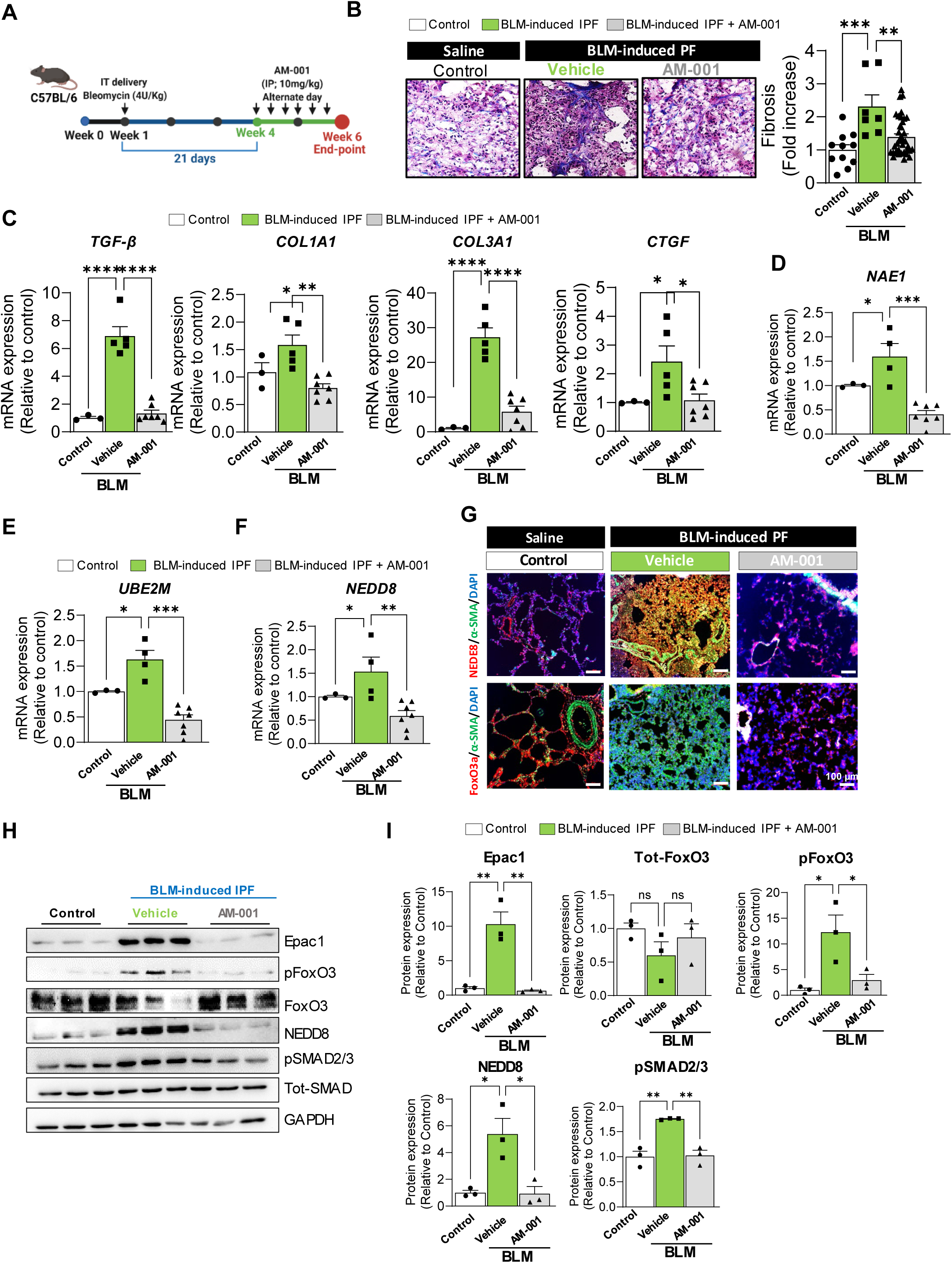
Pharmacological inhibition of Epac1 by AM-001 inhibits BLM-induced lung fibrosis. **A.** Schematic outlining the experimental design. C57BL/6 mice received a single intratracheal delivery of BLM (4 U/kg). After 21 days, AM-001 was administered via intraperitoneal injection every other day at a dose of 10 mg/kg for an additional 2 weeks. **B.** Masson’s trichrome staining of lung sections from control and BLM-challenged mice treated with either vehicle or AM-001. Representative images are presented in the left panel. The right panel shows the quantification of the percentage of the fibrotic area, expressed as the fold change relative to the saline control. **C.** Pro-fibrosis markers TGF-β, COL1A1, COL3A1, and CTGF mRNA expression levels in lung tissues from WT and Epac1^-/-^ mice treated with saline (vehicle) were measured by qRT-PCR in lung tissue from control and BLM-challenged mice treated with either vehicle or AM-001. **D-F.** The transcript levels of NAE1 (**D**), UBE2M (**E**) and NEDD8 (**F**) in lung tissues from both controls and BLM-challenged mice treated with either vehicle or AM-001 were analyzed by RT-qPCR. **G**. Co-immunostaining of NEDD8 (red), FoxO3a (red), and α-SMA (green) in lung sections from BLM-challenged mice treated with either vehicle or AM-001. Nuclei are stained in blue with DAPI. Scale bar-100 μm**. H.** Representative immunoblot analysis and (**I**) quantification of the indicated proteins in lung homogenates from the indicated group of mice. The data are presented as the mean ± SEM; *p < 0.05, ***p < 0.001, **p < 0.01.

### AM-001 as a modulator of immune lung cells in BLM-treated animals

To decipher the mechanism underlying the inhibition of BLM-induced lung fibrosis mediated by AM-001, we performed spectral flow cytometry to analyze immune cell populations in the lung of control and BLM-treated mice following the administration of AM-001. We evaluated the main leukocyte populations, including myeloid and T cells. This analysis revealed a distinct pattern of immune cell composition between BLM-induced fibrotic and AM-001 non-fibrotic mice (**Figure 7**).

**Figure 7.**
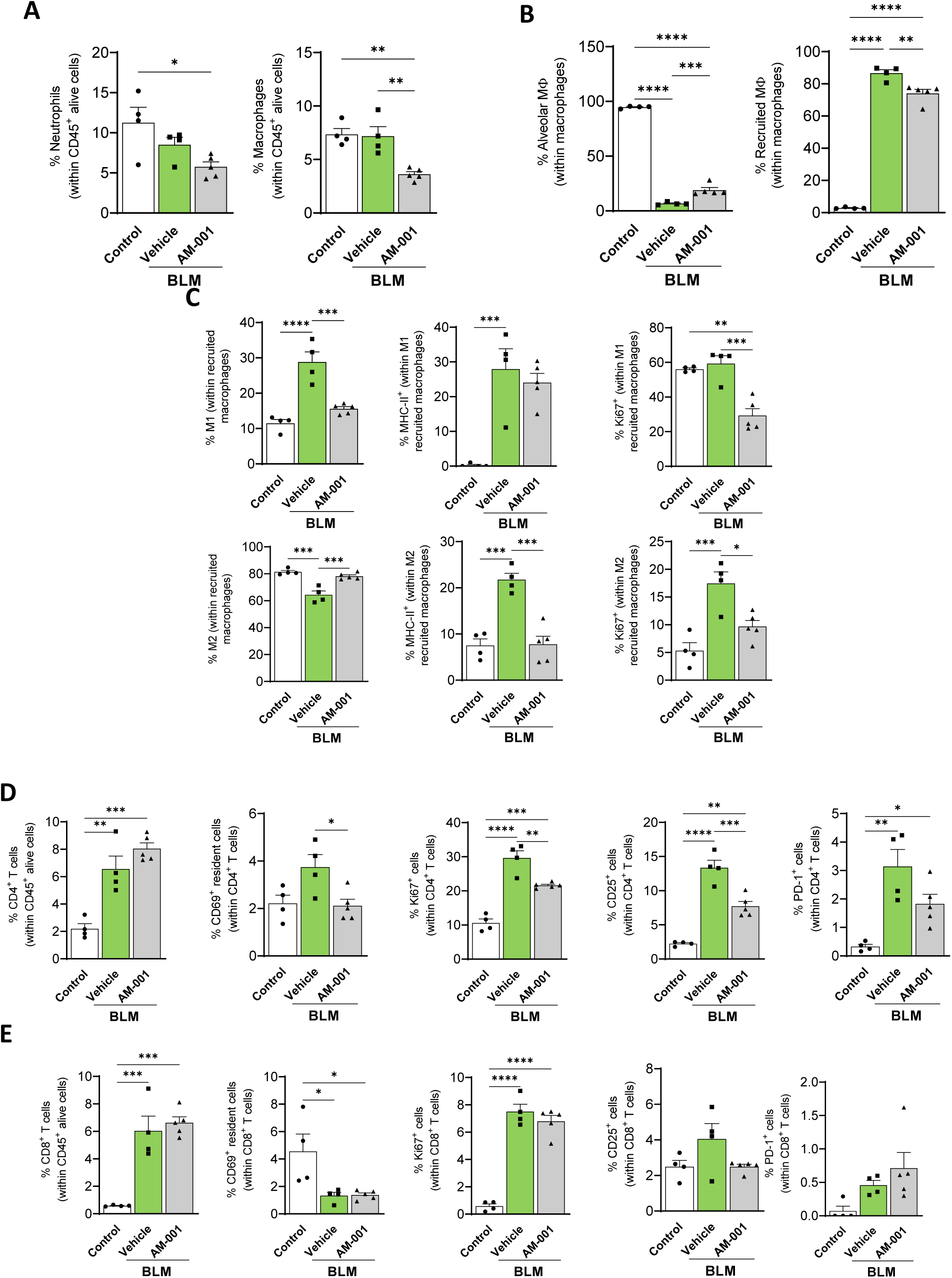
Phenotypical and functional characterization of immune responses induced by AM-001. C57BL/6 mice received either a single intratracheal delivery of BLM of 4 U/kg (vehicle) or left untreated (control). After 21 days, bleomycin-treated mice received AM-001 via intraperitoneal injections every alternate day at a dose of 10 mg/kg for an additional two weeks (AM-001). **A.** Percentage of neutrophils (Ly6G^+^CD3^-^CD19^-^) and macrophages (CD11c^+^F4/80^+^Ly6G^-^CD3^-^CD19^-^) within CD45^+^ alive cells. **B.** Percentage of alveolar (SiglecF^+^) and recruited (SiglecF^-^) macrophages (Mφ) within the total population. **C.** Percentage of M1 (Ly6C^+^, top) and M2 (Ly6C^-^, bottom) subsets within recruited macrophages, along with their expression of MHC-II and Ki67 within each subset. **D.** Percentage of CD4^+^ T cells (CD4^+^CD3^+^CD19^-^) within CD45^+^ alive cells, CD69^+^ resident CD4^+^ T cells, and expression of Ki67, CD25, and PD-1 within CD4^+^ T cell subset. **E.** Percentage of CD8^+^ T cells (CD8^+^CD3^+^CD19^-^) within CD45^+^ alive cells, CD69^+^ resident CD8^+^ T cells, and expression of Ki67, CD25, and PD-1 within CD8^+^ T cell subset. Results were analyzed using ordinary One-way ANOVA and Turkey’s multiple comparisons test *p ≤ 0.05, **p ≤ 0.01, ***p ≤ 0.001, **** p ≤ 0.0001.

The immune cell composition of the lungs showed a decrease in the percentage of neutrophils and macrophages following AM-001 treatment (**Figure 7A**). This is consistent with previous findings suggesting that the depletion of macrophages using clodronate-loaded liposomes in BLM-induced PF in mice suppressed the pathology of the disease^51^. Looking into the distinct macrophage subsets in the lungs, we observed that, while alveolar macrophages were markedly reduced in BLM-treated mice independent of the AM-001 administration, there was a marked increase in the number of recruited macrophages (**Figure 7B)**. Interestingly, the number of recruited macrophages was significantly decreased in the AM-001 groups, indicating that treatment 3 weeks after the induction of PF had a marked effect on lung infiltration by monocyte-derived cells. Further characterization of recruited macrophages showed that BLM-induced fibrotic mice exhibit significantly higher percentages of inflammatory M1 macrophages, while AM-001 treated mice display a regulatory M2 phenotype (**Figure 7C**). In addition, we observed a significant upregulation in the expression of MHC-II in M2 macrophages from fibrotic mice, facilitating the interaction with T cells and promoting cytokine secretion. Functionally, M1 and M2 recruited macrophages of the AM-001 treated group expressed lower percentages of the Ki67 proliferation marker, suggesting that mechanistically, AM-001 prevents PF in part by limiting the proliferative capacity of lung infiltrating monocytes (**Figure 7C**).

Characterization of the T cell-mediated immune response in the lungs revealed a significant increase in the percentages of both CD4^+^ and CD8^+^ T cells in the lungs of BLM-treated mice (**Figure 7D-E**). However, tissue-resident CD4^+^ T cells were significantly reduced in the AM-001 treated group with similar cell percentages as control non-fibrotic mice. Further evaluation of the T cell function indicated an overall reduction of the activation marker CD25 that achieved significant differences in CD4^+^ T cells (**Figure 7D-E**). This suggests that AM-001 treatment modulates the T cell response by reducing the ability of CD4^+^ T cells to respond to effector cytokines. Since CD25 controls T cell proliferation and differentiation in T cells, we next evaluated the expression of the Ki67 proliferation marker. Our results indicate that AM-001 treatment significantly reduced the proliferative capacity of CD4^+^ T cells. Further phenotypic characterization revealed the upregulation of programmed cell death protein 1 (PD-1) by CD4^+^ T cells in the lungs of BLM-treated mice (**Figure 7D-E**). PD-1 is an inhibitory receptor expressed by T cells during activation, which is associated with fibrotic lung disease in humans, and its upregulation in CD4^+^ T cells has been associated with PF following BLM administration in mice^52^.

### AM-001 treatment reverses fibrosis in *ex vivo* IPF human derived-PCLS model

Human precision-cut lung slices (PCLSs), three-dimensional (3D) tissue explants, provide a representative *ex vivo* platform model with preserved tissue lung tissue architecture and viability in culture ^53^. Therefore, we evaluated the potential therapeutic effects of AM-001 in human IPF disease mechanisms, Epac1 downstream signaling, and the expression of fibrotic markers in IPF-PCLS (**Figure 8A**). IPF human-derived-PCLS remained viable after 12 days of AM-001 treatment, and the fibrotic lesions confirmed by Masson Trichrome staining and extensive deposition of ECM were significantly diminished in IPF human-derived PCLS treated with AM-001 compared to vehicle (**Figure 8B-C**). Additionally, AM-001 reduced NEDD8 protein expression (red) in IPF human derived-PLCS co-stained with α-SMA (green), while FoxO3a (red) was restored (**Figure 8D-E**). Subsequently, IPF human-derived PCLS treated with AM-001 showed lower protein levels of fibrotic protein markers α-SMA, fibronectin, MMP-2, COL1, COL3, and total SMAD2/3, and trend reduction of NEDD8 and p-FoxO3a, and higher total FoxO3a protein levels compared to vehicle-treated IPF human derived-PCLSs (**Figure 8F-G**).

**Figure 8.**
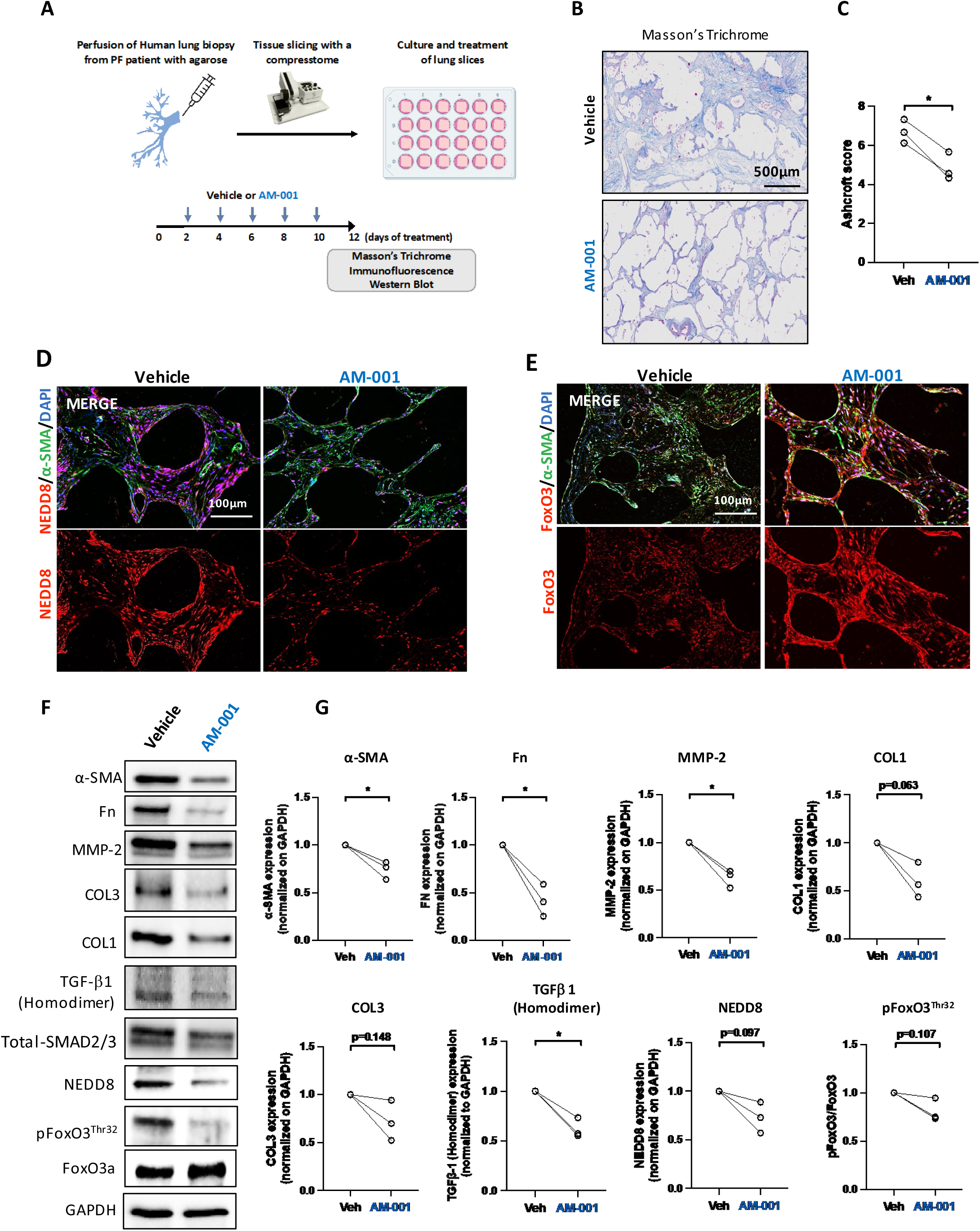
Pharmacological inhibition of Epac1 by AM-001 inhibits fibrosis in human-derived *ex vivo* precision-cut lung slices (PCLS) model of human lung fibrosis. **A.** IPF human-derived PCLS preparation and treatment with AM-001 every other day with 20μM AM-001 or vehicle for 12 days. **B**. Representative images of collagen fibers measured with Masson Trichrome staining and (**C**) quantified in IPF-PCLS treated AM-001 or vehicle as a control (*n* = 3). Scale bar-500 μm. **D-E**. Co-immunostaining of FoxO3a (red) and NEDD8 (red) with α-SMA in IPF**-**PCLS treated with vehicle or AM-001. Nuclei were counterstained with DAPI (blue). **F-G.** α-SMA, fibronectin (Fn), MMP-2, CoL1 and CoL-3, TGF-β, NEDD8, phospho- and total FoxO3a protein level were assessed by Western blot (WB) in IPF human derived**-**PCLS treated with AM-001 or the vehicle as a control. The data are presented as the mean ± SEM; *p < 0.05.

## Discussion

The current paradigm of pathological remodeling in PF by the dedifferentiation of fibroblasts to myofibroblasts and matrix production and deposition is driven by increased TGF-β levels in the lungs^18–20^. The precise role of Epac1 in PF and its potential as a therapeutic target has not been conclusively determined. This ambiguity arises from the predominance of *in vitro* studies that have yielded controversial results, which are likely influenced by the nature of the stimuli ^29–31^.

However, our data indicated that Epac1 plays a crucial role in profibrotic gene expression, fibroblast activation, and proliferation in PF, suggesting that Epac1 could be a viable target for developing AM-001 as a novel anti-fibrotic agent. Moreover, accumulating evidence underscores Epac1 as a key effector in the lung, notably in modulating airway inflammation and proliferation^54–32^. Huang et al. demonstrated that while prostaglandin-mediated inhibition of collagen expression in primary human fetal lung fibroblasts was exclusively mediated by PKA, Epac1 was responsible for inhibiting fibroblast proliferation^55^. However, there are some controversies regarding the role of Epac1 in cell proliferation, migration^56^, and fibrosis^55–30^. Depending on the cell type and its differentiation state, some studies have reported an inhibitory role^57,29^. Additionally, other studies have shown opposite results^58^; thus, Epac1 appears to be critical for the integration of pro- and anti-fibrotic signals and the regulation of fibroblast function^30^. Interestingly, Epac1 and PKA mediate the opposing effects of cAMP on PKB/AKT regulation, which may provide a potential mechanism for the differential cell type-specific effects of cAMP^59^. Therefore, the apparent cellular effects of cAMP may vary depending on the relative cellular abundance and compartmentation of Epac1 and PKA, as well as stress inducers. Several studies have suggested that Epac1 is a promising target for the treatment of various disorders. Compelling evidence indicate that Epac1 inhibition may be an effective approach for treating cancer and cardiovascular diseases ^60^.

Importantly, Epac1 KO mice exhibit decreased fibrosis and improved cardiac function under stress conditions such as chronic isoproterenol infusion. In line, we showed here that Epac1 KO mice were protected against the BLM-induced PF phenotype. Therefore, the identification of pharmacological modulators is important for unraveling the role of Epac1 in disease conditions ^60^.

AM-001 selectively inhibited Epac1 catalytic activity in cultured cells in response to the selective cAMP analog 8-pCPT^58^. AM-001 also exerts cardioprotective effects against myocardial ischemia/reperfusion injury and prevents chronic β-adrenergic (AR) activation, induced cardiac inflammation, and fibrosis^58^. Nonetheless, the precise role of AM-001 Epac1 inhibitor in lung diseases and PF has not been explored and remains unclear.

Mechanistically, AM-001 targets key pathways involved in IPF, such as the TGF-β-SMAD2/3, IL-6/STAT3, and phospho-AKT/phospho-FoxO3a signaling pathways. We found that Epac1 expression was increased in BLM-treated WT mice and that AM-001 treatment reduced Epac1/TGF-β, collagen, and IL-6 gene expression and signaling, which could be related to FoxO3a pathway activation and TGF-β signaling suppression, thus reversing lung fibrosis. FoxO3a has been shown to play a key role in fibrogenesis, and its expression was reduced in IPF^34^. FoxO3a, a target of PKB/AKT via phosphorylation, ubiquitination, and degradation, has been implicated in the differentiation and hyperproliferation of lung FB ^34, 39^. Additionally, FoxO3a KO mice displayed enhanced susceptibility to BLM and increased fibrosis and mortality^34^. We validated this result and showed that FoxO3a expression was downregulated in IPF patients and animal models of PF, while FoxO3a phosphorylation was increased.

Neddylation is a reversible and important post-translational modification that involves the conjugation of the ubiquitin-like protein neuronal precursor cell-expressed developmentally downregulated protein 8 (NEDD8) to target protein degradation ^64^. This process involves NEDD8-activating enzyme (NAE, a heterodimer comprising subunits NAE1 and UBA3, ubiquitin-activating enzyme 3), NEDD8-conjugating enzyme E2 (UBC12), and substrate-specific E3s ^65–67^. NEDD8 is first adenylated and activated by E1 NEDD8-activating enzyme (NAE), a heterodimer consisting of NAE1 and UBA3. NEDD8-loaded NAE is transferred to one of the two NEDD8-conjugating enzymes, UBE2M (also known as UBC12) or UBE2F, through a trans-thiolation reaction ^68, 69^. However, the effects of NEDD8-mediated neddylation and degradation of FoxO3A and its function in PF, remain unclear. Notably, the expression and phosphorylation of FoxO3a by Epac1 via AKT activation were inhibited by AM-001 and the NEDD8 inhibitor MLN4924. These findings suggest that NEDD8 is involved in FoxO3a degradation.

NEDD8 targets proteins and affects their stability, subcellular localization, conformation, and function^66^. It has been reported that FoxO3a phosphorylation is necessary for its nuclear localization^70^. Here, we showed that NEDD8 triggers phospho-FoxO3a degradation and the subsequent proteasomal degradation of FoxO3a. However, MLN4924 suppressed the expression of profibrotic markers, and the combination of MLN4924 and AM-001 potentiated this effect. These data showed that the pharmacological inhibition of Epac1 and, consequently, the neddylation pathway suppressed IPF-FB activation and the PF phenotype.

Resident myeloid cells continuously sample the tissue microenvironment that triggers inflammatory responses to tissue injury and trauma and pave the way for the progressive recruitment of circulating myeloid cells, such as neutrophils and inflammatory macrophages^71^, and represent a key factor in many diseases, including IPF^72, 73, 74, 75, 76^. Epac1 is expressed in immune and hematopoietic cells, and its activation promotes monocyte adhesion and polarization and enhances chemokine-induced migration^77^. Our results indicate that Epac1 inhibition with AM-001 decreased neutrophil and macrophage infiltration in the lungs, which is associated with a reduced pathology in BLM-induced lung fibrosis^78^. Interestingly, AM-001 treatment also reduced the number of Ly6Chi monocyte-derived M1 macrophages that facilitate the progression of PF^51^. BLM-induced pathogenic lung macrophages also express higher levels of MHC-II and exhibit increased proliferation rates^79^, which induce cytokine production in CD4+ T cells^80^. Epac1 also regulates TGF-β1 signaling in CD4^+^ T cells by regulating p-STAT3 and SMAD4^81^ in a subset of PD-1 expressing CD4^+^ T cells^52^. Our data indicate that AM-001 treatment reduced the number of CD4^+^PD-1^+^ T cells in the lungs. The anti-inflammatory properties of Epac1 inhibitors further emphasize the importance of Epac1 inhibition as a promising new therapeutic target for the treatment of respiratory diseases associated with excessive and persistent inflammatory responses ^20, 22, 82^.

Finally, for translational novel therapy and to closely mimic *in vivo* PF conditions and evaluate the effects of AM-00, we applied disease models to human-derived IPF-PCLS that conserved the anatomical architecture of the lung tissue microenvironment of the fibrotic lung and ECM. In human-derived IPF-PCLS, AM-001 behaved as a therapeutic compound since it reduced fibrotic lesions, collagen deposition, and ECM remodeling, as well as myofibroblast activation. Of particular interest, NEDD8 and pFoxO3a were also decreased in AM-001 treated IPF human-derived PLCs compared to the control vehicle.

In conclusion, our findings demonstrated that pharmacological inhibition of Epac1 using AM-001 suppressed the fibrotic characteristics of IPF-FBs *in vitro*, experimental lung fibrosis *in vivo*, and in a human-derived IPF-PCLS model *ex vivo*, thus providing strong evidence that targeting Epac1 pathway is a potential new therapy for IPF.

## Author Contributions

MB, LH, SB, and FL contributed to the experimental design. KJ, MB, SE, DLO, LPR, KF, VJ, LH, and SZ were responsible for data acquisition. Resources were provided by MB, LH, FL, JM, JC, SB, and J.O. Data analysis was performed by KJ, MB, SE, DLO, LPR, KF, VJ, and LH. All authors participated in manuscript writing, reviewing, and editing. All authors have read and agreed to the published version of the manuscript.

## Data Availability Statement

All data generated or analyzed during this study are included in this published article and its Supplementary Information Files. Any additional datasets used and/or analyzed during the current study are available from the corresponding author upon reasonable request.

## Abbreviations

AKT: Protein Kinase B
AR: Adrenergic Receptor
BLM: Bleomycin
cAMP: Cyclic Adenosine Monophosphate
CTGF: Connective Tissue Growth Factor
CEIC: Committee for Ethics in Clinical Research
COL1A1: Collagen Type I Alpha 1
COL3A1: Collagen Type III Alpha 1
CTL: Control
DEGs: Differentially Expressed Genes
ECM: Extracellular Matrix
Epac: Exchange Proteins Directly Activated by cAMP
FBS: Fetal Bovine Serum
FGM-2: Fibroblast Growth Medium-2
FoxO3a: Forkhead Box O3
GEF: Guanine Nucleotide Exchange Factor
GO: Gene Ontology
IL-6: Interleukin-6
IPF: Idiopathic Pulmonary Fibrosis
KO: Knock-out
MLN4924: Pevonedistat, NEDD8 activating enzyme inhibitor
NAE: NEDD8-Activating Enzyme
NHL-FB: Normal Human Lung Fibroblasts
pAKT: Phosphorylated Protein Kinase B
PF: Pulmonary Fibrosis
RAPGEF3: RAP Guanine Nucleotide Exchange Factor 3
RIN: RNA Integrity Number
shRNA: Short Hairpin RNA
TGF-β: Transforming Growth Factor Beta
TGF-β1: Transforming Growth Factor Beta 1
TNC: Tenascin C
WT: Wild-Type

---

**Supplementary Figure 1. Epac isoforms expression levels in IPF and BLM-challenged mice**. **A.** Epac2 mRNA level in lung tissue from healthy donors and IPF patients. **B**. Quantification of αSMA and CyclinD1 protein expression levels normalized to those of GAPDH in isolated primary IPF-FBs (*n* = 3) and NHL-FB cells (*n* = 3). **C**. Epac1 mRNA in lung tissues from sham control vehicle-injected mice and BLM-challenged mice analyzed by qRT-PCR *(n*= 3). **D**. Representative Immunoblot analysis and quantification of Epac1 protein levels were normalized to those of GAPDH. The data are presented as the mean ± SEM; *p < 0.05, **p ≤ 0.01, ***p < 0.001, ns: not significant.

**Supplementary Figure 2. Epac isoforms levels in NHL-FBs. A.** Epac1 mRNA expression level in NHL-FBs overexpressing a control GFP adenovirus (Ad.CT) or Epac1 adenovirus (Ad.Epac1) for 48 h. **B.** Epac2 mRNA expression in NHL-FBs overexpressing Epac1(Ad.Epac1) measured by qRT-PCR. **C.** Epac2 mRNA expression in normal NHL-FBs overexpressing a non-silencing shRNA (shNS) or a specific shRNA against Epac1 (sh.Epac1) measured by qRT-PCR. Data are presented as the mean ± SEM; ***p < 0.001, ns: not significant.

**Supplementary Figure 3. Effects of CE3F4 on normal human lung fibroblasts. A.** Chemical structure of AM-001 a thieno [2,3-b]pyridine scaffold composed of three structural units containing thiophenyl, phenyl, and fluorophenylamide moieties that specifically targets Epac1 activity. **B**. Chemical structure of the equine inhibitor CE3F4 that targets Epac1 activity. **C.** Experimental design for *in vitro* experiments. Briefly, NHL-FBs were treated with 20 µM CE3F4 for 24 hours followed by treatment with TGF-β for an additional 24 hours. **D**. Transcript levels of the profibrotic genes in NHL-FBs treated as described in panel B and measured by qRT-PCR. The data are presented as the mean ± SEM; *p < 0.05, **p < 0.01, ***p < 0.001.

**Supplementary Figure 4. Effects of Epac1 knock-down on FOXO3 levels in NHL-FBs and IPF-FBs. A.** FOXO3 mRNA expression in NHL-FBs overexpressing Ad.CT or Epac1 (Ad.Epac1) measured by qRT-PCR**. B.** mRNA expression levels of FoxO3 in NHL-FBs or IPF-FBs overexpressing either shNS or sh.Epac1. **p < 0.01, ***p < 0.001.

**Supplementary Figure 5. Effects of AM-001 and Epac1 knock-down on the neddylation pathway in NHL-FBs and IPF-FBs. A.** Epac1 mRNA expression in nonactivated NHL-FBs or IPF-FBs treated with AM-001 or DMSO (as a vehicle control) for 48 hours. **B**. UBE2F and UBA3 mRNA expression in NHL-FBs treated with AM-001 or DMSO (as a vehicle control) for 48 hours. **C-G**. NAE1, NEDD8, UBE2M, UBE2F and UBA3 mRNA expression in NHLF or IPF-FBs overexpressing either shNS or shEpac1 in nonactivated NHLF-FB and IPF-FB. The data are presented as the mean ± SEM; *p < 0.01, **p < 0.01, ***p < 0.001. ns: not significant.

**Supplementary Figure 6. mRNA levels of FGFR1 and TNC in lung tissues from IPF patients and healthy controls. A.** FGFR1 mRNA levels were assessed in lung tissues from IPF patients and healthy control lungs (n = 4-8) using qRT-PCR. **B.** TNC mRNA levels were assessed in lung tissues from IPF patients and healthy control lungs (n = 4-8) using qRT-PCR. The data are presented as the mean ± SEM; *p < 0.05, ***p < 0.001.

**Supplementary Figure 7. Pharmacological inhibition and Epac1 knock-down decreased FGFR1 and TNC mRNA expression in NHL-FBs and IPF-FBs. A-B.** FGFR1 and TNC mRNA expression were measured by qRT-PCR in NHL-FBs and IPF-FBs treated with AM-001 or DMSO (as a vehicle control) for 48 hours. **C-D.** FGFR1 and TNC mRNA expression were measured by qRT-PCR in NHL-FBs and IPF-FBs expressing a non-silencing shRNA (shNS) or a specific shRNA against Epac1 (shEpac1) for 72 hours. The data are presented as the mean ± SEM; *p < 0.05, **p < 0.01, ***p < 0.001.

**Supplementary Figure 8. Epac1, Epac2, FGFR1, and TNC expression levels in lung tissues from WT and Epac1 KO mice post-BLM challenge. A.** Epac1 mRNA expression in lung tissues from wild-type (WT) and Epac1 knock-out (Epac1^-/-^) mice subjected to a single intratracheal aerosolized delivery of bleomycin (BLM, 4 U/kg) or vehicle (saline). **B.** Epac2 mRNA expression in lung tissues from WT and Epac1^-/-^ mice following treatment with BLM or vehicle. **C.** FGFR1 mRNA expression in lung tissues from WT and Epac1^-/-^ mice challenged with BLM or vehicle. **D.** TNC mRNA expression in lung tissues from WT and Epac1^-/-^ mice after BLM or vehicle administration. The mRNA expression levels were assessed by RT-qPCR and the data are presented as the mean ± SEM; *p < 0.05, **p < 0.01, ***p < 0.001. ns: not significant.

**Supplementary Figure 9. Expression of FGFR1 and TNC in lung tissues from C57BL/6 mice treated with AM-001 following BLM challenge. A-B.** mRNA expression of FGFR1 and TNC by qRT-PCR in lung tissues from mice treated with AM-001 or vehicle following BLM challenge. The data are presented as the mean ± SEM; *p < 0.05, **p < 0.01, ***p < 0.001. ns: not significant.

**Supplementary Figure 10. Vascular remodeling in lung sections from C57BL/6 mice treated with AM-001 following BLM challenge. A.** Vascular remodeling assessed by hematoxylin and eosin (H&E) staining of lung sections of indicated mice. Representative images of small pulmonary arteries are provided. **B.** Quantification of media wall thickness in these arteries is shown as fold change. The data are presented as the mean ± SEM; ***p < 0.001.

**Supplementary Figure 11. Cardiac hemodynamics and remodeling in C57BL/6 mice treated with AM-001 following BLM challenge. A.** Right ventricular systolic pressure (RVSP) is measured by right heart catheterization during open-chest surgery using a pressure catheter directly inserted into the RV. **B.** Fulton Index, defined as the ratio of the RV weight to the LV weight plus septum weight, is used to measure RV hypertrophy levels. **C.** mRNA levels of ANP, BNP, and β-MHC in RV samples from these mice, were measured by qRT-PCR. **D.** Representative images of RV fibrosis assessed by Masson’s Trichrome staining. **E.** Quantification of fibrosis using Masson’s trichrome staining, with the blue-stained area measured and expressed as fold change for comparison. **F-G.** Representative images and quantification of right ventricular (RV) cardiomyocyte size were assessed using wheat germ agglutinin (WGA) staining. The data are presented as the mean ± SEM; *p < 0.05, **p < 0.01, ***p < 0.001.

