## Supplementary Figures for "Pharmacological Inhibition of Epac1 Protects against Pulmonary Fibrosis by Blocking FoxO3a Neddylation"

### Supplementary Figure 1

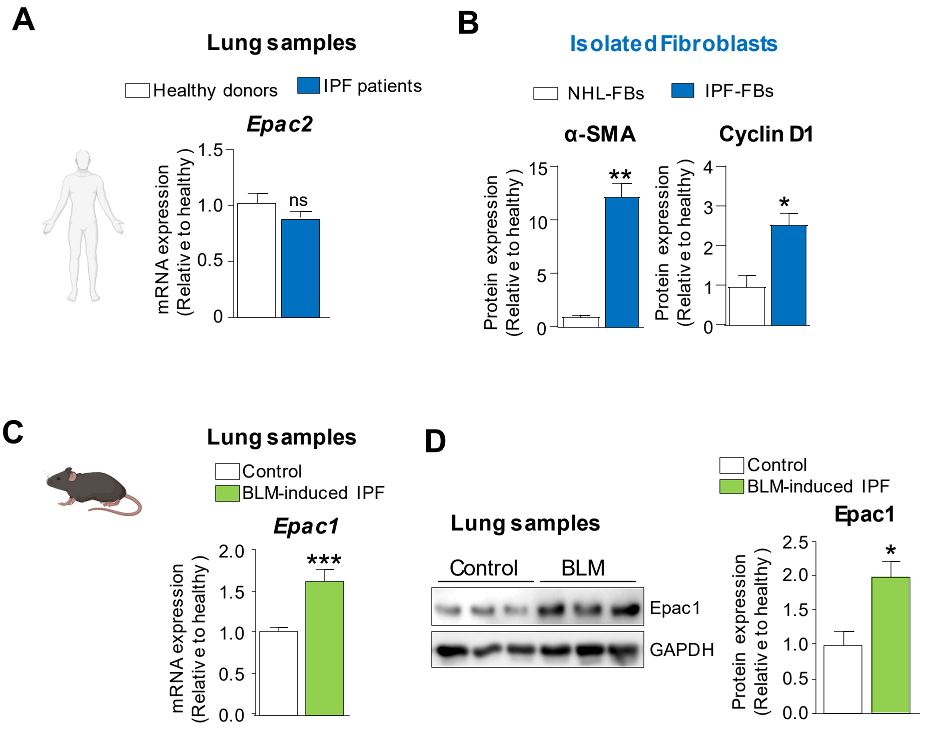

### Supplementary Figure 2

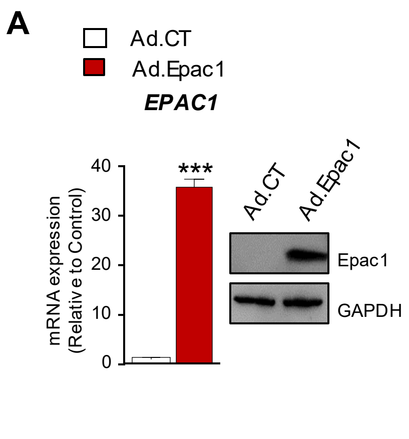

**Supplementary Figure 3**

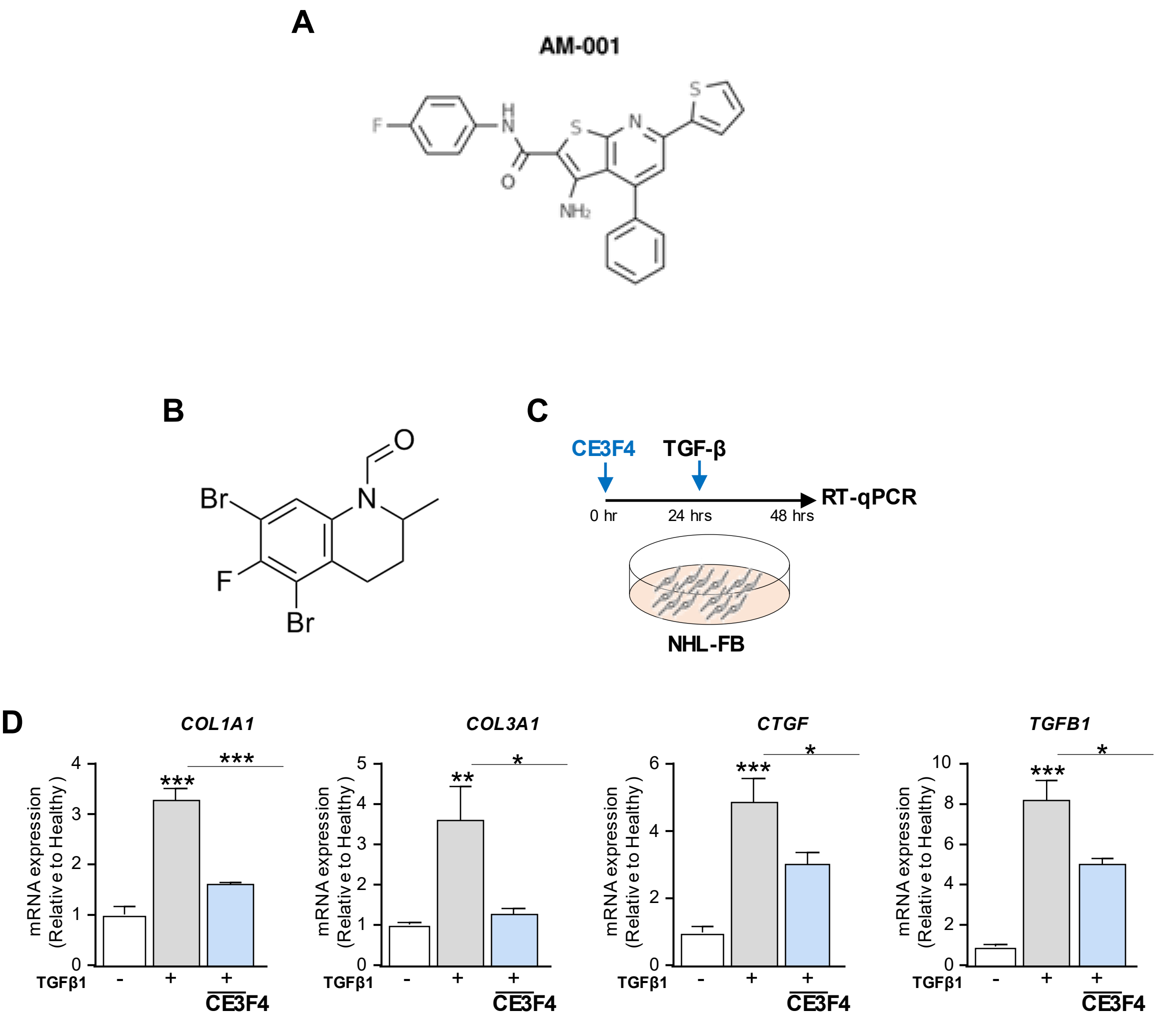

### Supplementary Figure 4

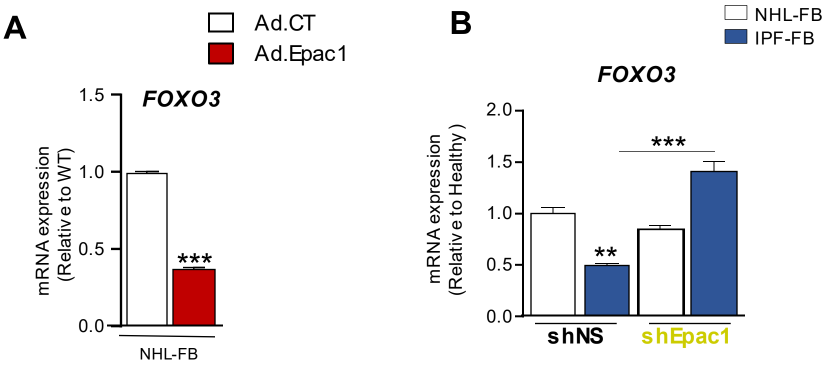

### Supplementary Figure 5

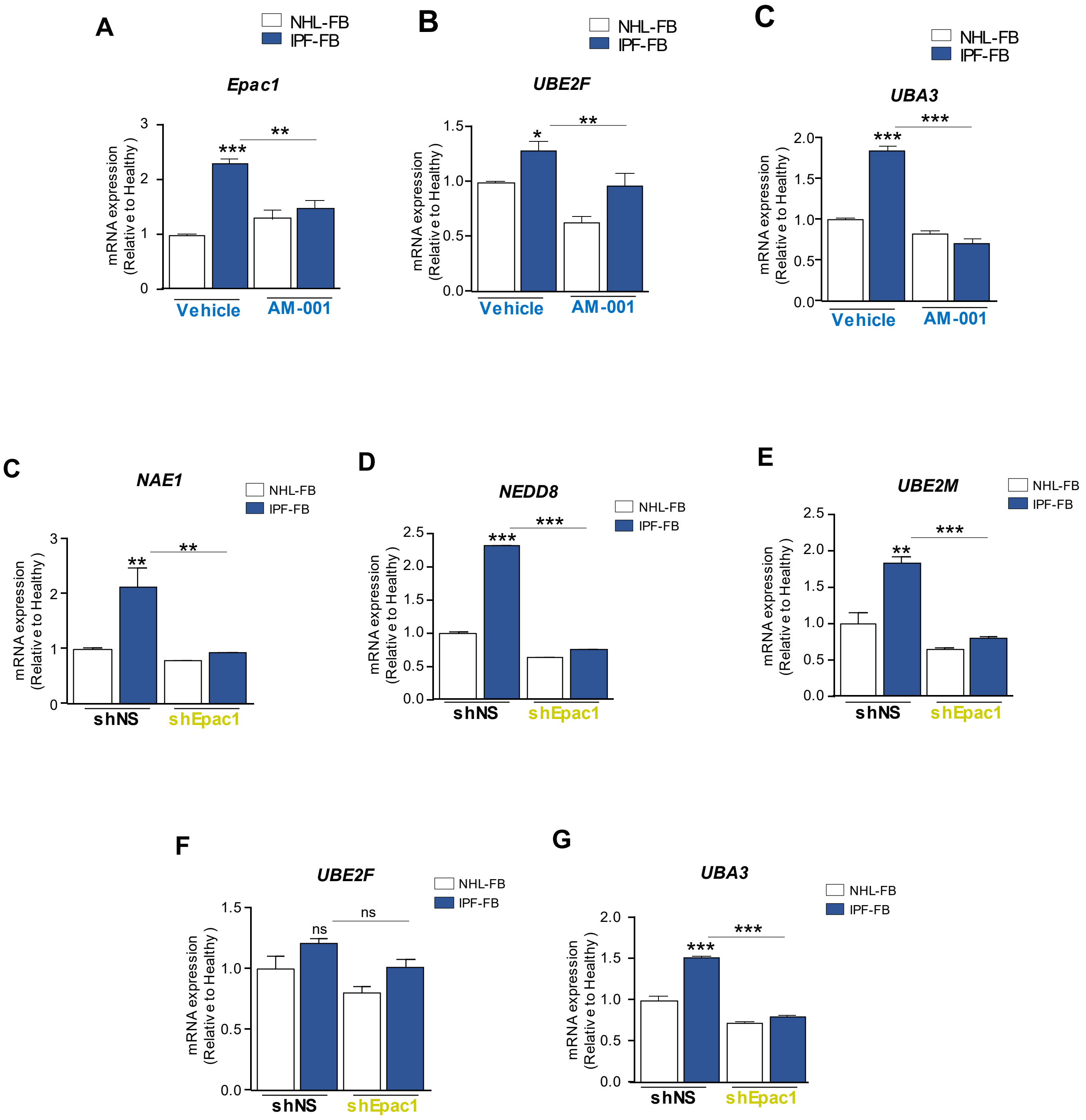

### Supplementary Figure 6

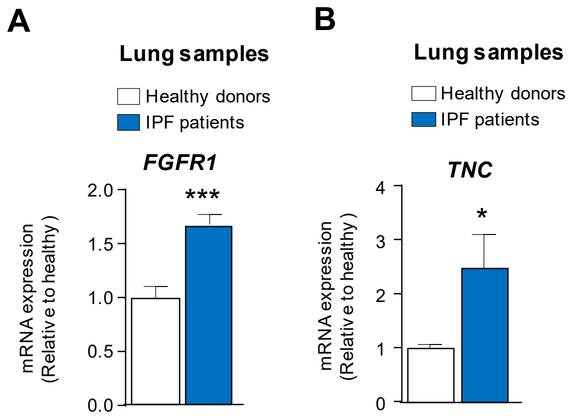

### Supplementary Figure 7

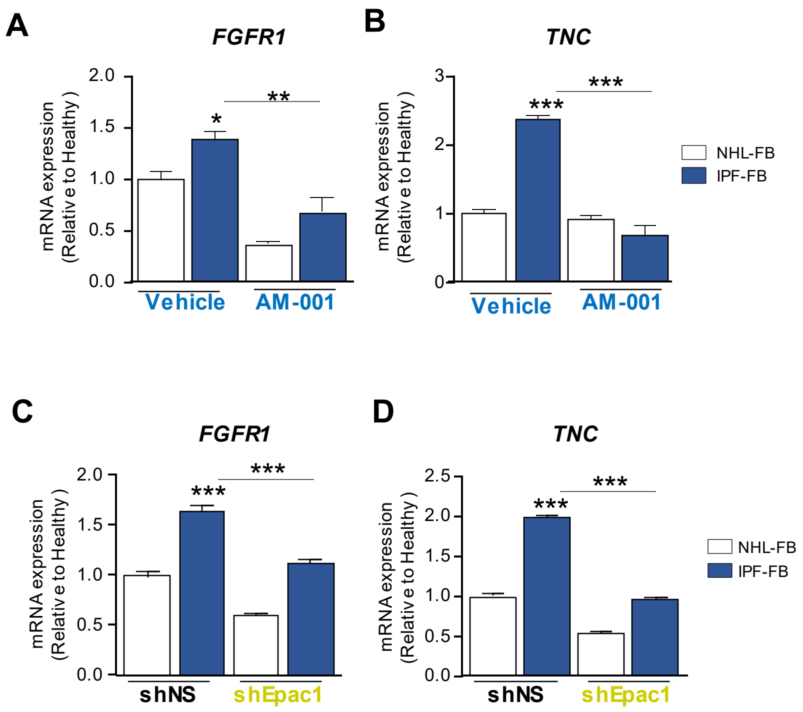

### Supplementary Figure 8

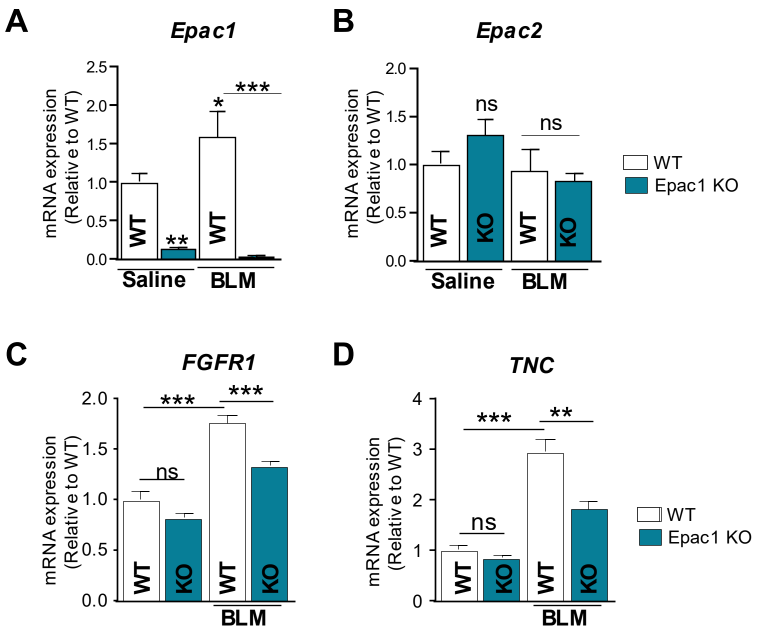

### Supplementary Figure 9

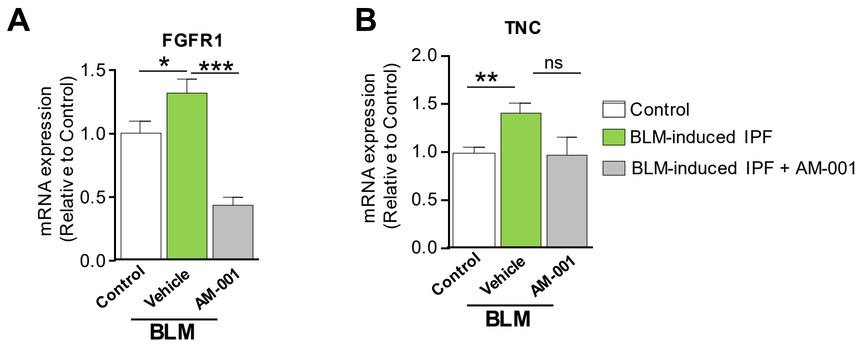

#### Supplementary Figure 10

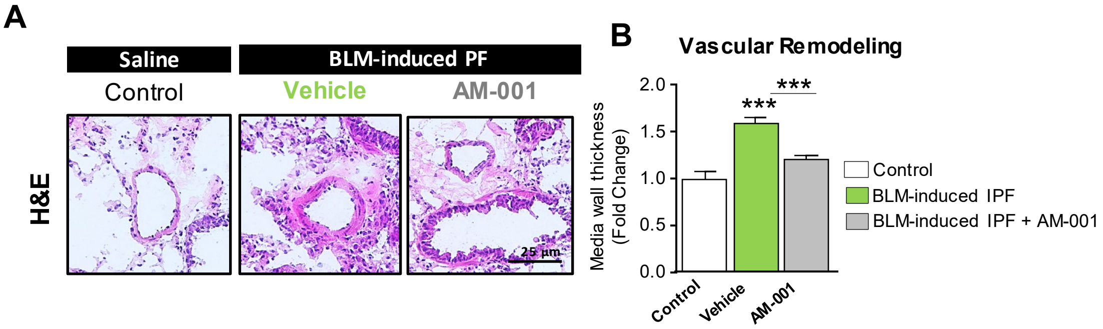

### Supplementary Figure 11

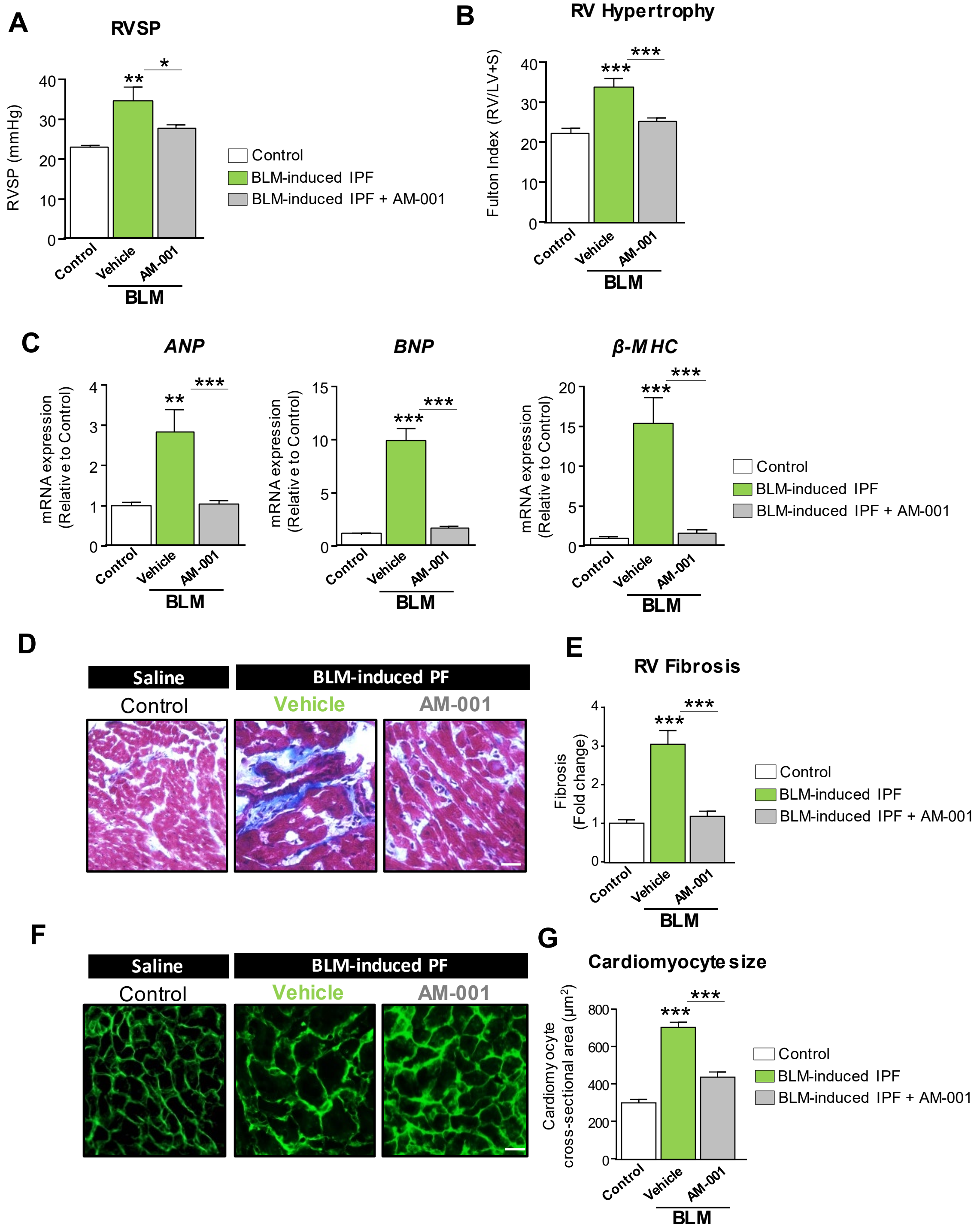
