## Supplementary Tables for "Pharmacological Inhibition of Epac1 Protects against Pulmonary Fibrosis by Blocking FoxO3a Neddylation"

### Supplement Materials

#### Supplementary Table 1. IPF Patients Clinical Data

| ID | Age/Sex | Diagnosis | FIV1 (%pred) | FVC (%pred) | FIV1/FVC | DLco (% Diffusion capacity CO) | (TLC) partial pressure of oxygen (mmHg) | partial pressure of CO2 (mmHg) | Alveolar-arterial gradient | PaO2 | 6 minutes walking test (6MWT; meters) | N° of reagudizations/year |
|---|---|---|---|---|---|---|---|---|---|---|---|---|
| Patient 1 | 71/MAN | IPF | 72 | 64 | 83.03 | 49 | 76.8 | 36.7 | 28.3 | 95.9 | 606 | 0 |
| Patient 2 | 72/MAN | IPF | 54 | 56 | 70 | 29 | 58.7 | 41 | 40 | 93 | 364 | 0 |
| Patient 3 | 60/MAN | IPF | 50 | 45 | 86 | 35 | 65.7 | 35.4 | 40.5 | 90 | 552 | 0 |
| Patient 4 | 60/MAN | IPF | 36 | 32 | 87 | 21 | 66 | 32 | 42.7 | 91 | 390 | 1 |
| Patient 5 | 68/MAN | IPF | 70 | 61 | 84 | 40 | 83 | 36 | 22 | 84 | 557 | 0 |
| Patient 6 | 65/MAN | IPF | 62 | 61 | 83 | 46 | 75 | 36 | 36 | 95 | 542 | 0 |
| Patient 7 | 68/MAN | IPF | 59 | 35 | 60 | 26 | 54 | 42 | 26 | 92 | 368 | 1 |
| Patient 8 | 71/MAN | IPF | 71 | 45 | 64 | 36 | 69 | 32 | 37 | 93 | 458 | 0 |

#### Supplementary Table 2. Control Healthy Non-IPF Subjects

| ID | Age/Sex | Diagnosis | FIV1 (%pred) | FVC (%pred) | FIV1/FVC | DLco (% Diffusion capacity CO) | (TLC) partial pressure of oxygen (mmHg) | partial pressure of CO2 (mmHg) | Alveolar-arterial gradient | PaO2 | 6 minutes walking test (6MWT) (meters) | N° of reagudizations/year |
|---|---|---|---|---|---|---|---|---|---|---|---|---|
| Patient 1 | 71/FEMALE | Healthy donors | ND | ND | ND | ND | ND | ND | ND | ND | ND | ND |
| Patient 2 | 70/MAN | Healthy donors | ND | ND | ND | ND | ND | ND | ND | ND | ND | ND |
| Patient 3 | 65/MAN | Healthy donors | ND | ND | ND | ND | ND | ND | ND | ND | ND | ND |
| Patient 4 | 54/MAN | Healthy donors | ND | ND | ND | ND | ND | ND | ND | ND | ND | ND |
| Patient 5 | 78/MAN | Healthy donors | ND | ND | ND | ND | ND | ND | ND | ND | ND | ND |
| Patient 6 | 64/MAN | Healthy donors | ND | ND | ND | ND | ND | ND | ND | ND | ND | ND |
| Patient 7 | 62/MAN | Healthy donors | ND | ND | ND | ND | ND | ND | ND | ND | ND | ND |
| Patient 8 | 71/MAN | Healthy donors | ND | ND | ND | ND | ND | ND | ND | ND | ND | ND |

### Supplementary Table 3

| ID | Category | Diagnosis | Age | Sex |
|---|---|---|---|---|
| **Patient 1** | Fibrosis | IPF-PH | 68 | Male |
| **Patient 2** | Fibrosis | PF | 58 | Male |
| **Patient 3** | Fibrosis | IPF | 77 | Female |

**Supplementary Table 4.** Primer sequences for RT-qPCR analysis; clone ID and catalog numbers for shRNAs and siRNA (Open Biosystems); antibodies used; source and concentration of chemical inhibitors used.

| **Application** | **Gene symbol** | **Species** | **Forward primer (5′-3′)** | **Reverse primer (5′-3′)** |
|---|---|---|---|---|
| **RT-qPCR** | Epac1 | Human Mouse | CTGCTCTTTGAACCACACAGCAAG | GGTTGAAGTCCTGCTTGTCCAC |
| | Epac2 | Human Mouse | AAACCACCTGGCCAGAGGACT | TTCCCCCTGGTTAAACAACACAGT |
| | COL1A1 | Mouse Human | TTCAGTGGTTTGGATGGTGCCAA | CCAGCTTCACCCTTAGCACCA |
| | COL3A1 | Mouse Human | GAGATGTCTGGAAGCCAGAACCATG | ATCTCCCTTGGGGCCTTGAGGT |
| | CTGF | Mouse Human | GTCTGCGCCAAGCAGCTG | ACGTCCATGCTGCACAGGG |
| | TGFB | Mouse Human | CCTGCAAGACCATCGACATGGAG | GGTCGCGGGTGCTGTTGTA |
| | SMA | Human | CACCAACTGGGACGACATGG | CCATCTCCAGAGTCCAGCAC |
| | NAE1 | Human | CCAGTAATAGAATCTCATCCAGATAATGCA | TAGCTATGATCACAATCCATGGAGTATG |
| | NEDD8 | Human Mouse | CGAATCAAGGAGCGTGTGGA | TCTTGTAATCAGCTGCTGTCTTCTC |
| | UBE2M | Human Mouse | AACATTGACCTCGAGGGCAAC | CAAGAAGAGATACTGCAGGCCATAA |
| | FOXO3 | Human Mouse | AATTCTGTCAGCAACATGGGCTTGA | GCTCCCATTGAACATGTCCAGGT |
| | UBA3 | Human | ATGTAAAGTTCTAGTCATTGGAGCT | ATTCTGCAGCAACTTCAGCC |
| | UBE2F | Human Mouse | GGTTACTACCAGGGTGGAAAAT | TGTGATGTTGGGGTGCCAG |
| | FGFR1 | Human Mouse | TGCAGAGCATCAACTGGCTG | ACATTGACGGAGAAGTAGGTGGT |
| | TNC | Human Mouse | ATGCAGTTCCGTGTGCCTG | ACTTCCACAGATGTCTCCTTGAT |
| | ANP | Mouse | GCTTCCAGGCCATATTGGAG | GGGGGCATGACCTCATCTT |
| | BNP | Mouse | CTGGGAAGTCCTAGCCAGTC | TTTTCTCTTATCAGCTCCAGCA |
| | β-MHC | Mouse | ACTGTCAACACTAAGAGGGTCA | TTGGATGATTTGATCTTCCAGGG |
| | IL6 | Human | AGCTGCAGGCACAGAACCAG | ATTTGCCGAAGAGCCCTCAG |
| | | Mouse | TGGGACTGATGCTGGTGACA | GCCTCCGACTTGTGAAGTGGT |
| | GAPDH | Human | CGACCACTTTGTCAAGCTCA | AGGGGAGATTCAGTGTGGTG |
| | GAPDH | Mouse | GTGAAGGTCGGTGTGAACG | TCGTTGATGGCAACAATCTC |
| shRNAs | **Gene symbol** | **Clone ID** | | **Catalog number** |
| | *RAPGEF3 (Epac1)* | TRCN0000047228 | | RHS3979-201771060 |

| Immunoblotting | **Protein symbol** | **Antibody source** | **Dilution** |
|---|---|---|---|
| | Epac1 | Cell signaling | 1:1000 |
| | Alpha-SMA | Sigma Aldrich | 1:1000 |
| | CyclinD1 | BD Pharmingen | 1:1000 |
| | pSTAT3 | Cell signaling | 1:1000 |
| | Tot-STAT3 | Cell signaling | 1:1000 |
| | NEDD8 | Cell signaling | 1:1000 |
| | pSMAD2-3 | Cell signaling | 1:1000 |
| | Tot-SMAD2-3 | Cell signaling | 1:1000 |
| | pAkt | Cell signaling | 1:1000 |
| | Tot-Akt | Cell signaling | 1:1000 |
| | $pFOXO3^{Thr32}$ | Cell signaling | 1:1000 |
| | Tot-FOXO3 | Cell signaling | 1:1000 |
| | Fibronectine | Abcam | 1:500 |
| | COL1 | ThermoFisher Scientific | 1:500 |
| | COL3 | Proteintech | 1:3000 |
| | MMP-2 | Cell signaling | 1:1000 |
| | GAPDH | Cell signaling | 1:5000 |
| Pharmacological agents | **Compounds** | **Concentration** | **Source** |
| | TGF-beta | 5 nM | Peprotech |
| | SB431542 | 10 µM | MedChem Express |
| | Bleomycin | 4U/Kg | MedChem Express |
| | MG132 | 10 µM | MedChem Express |
| | MLN4924 | 5 µM | MedChem Express |
| | AM-001 | 20 µM | Gift from Dr. Frank Lezoualch |
| | CE3F4 | 20 µM | Gift from Dr. Frank Lezoualch |
| | 8-CPT | 5 nM | Biolog |

**Supplementary Table 5. Antibodies used for immune phenotyping by flow cytometry.**

| Target | Conjugate | Clone | Company | Reference | Dilution |
|---|---|---|---|---|---|
| CD62L | BV421 | MEL-14 | BioLegend | 104436 | 1:100 |
| CD11b | SuperBright436 | M1/70 | Invitrogen | 62-0112-82 | 1:80 |
| Foxp3 | PB | MF-14 | BioLegend | 126410 | 1:120 |
| CD8 | BV510 | 53-6.7 | BioLegend | 100752 | 1:150 |
| CD19 | BV570 | 6D5 | BioLegend | 115535 | 1:150 |
| MHCII | BV605 | M5/114.15.2 | BioLegend | 107639 | 1:100 |
| NK.1.1 | BV650 | PK136 | BioLegend | 108736 | 1:60 |
| CD25 | BV711 | PC61 | BioLegend | 102049 | 1:120 |
| CD45 | BV750 | 30-F11 | BD Biosciences | 746947 | 1:150 |
| CD206 | BV785 | C068C2 | BioLegend | 141729 | 1:129 |
| CD4 | PerCP | GK1.5 | BioLegend | 100432 | 1:100 |
| CD11c | PerCP-Cy5.5 | N418 | BioLegend | 117328 | 1:75 |
| Ki67 | PerCP-eFluor710 | SolA15 | Invitrogen | 46-5698-80 | 1:200 |
| CD86 | PE | RMMP-2 | Invitrogen | MA5-17953 | 1:60 |

| | | | | | |
|---|---|---|---|---|---|
| NKp46 | PE-Dazzle594 | 29A1.4 | BioLegend | 137630 | 1:60 |
| CD69 | PE-Cy5 | H1.2F3 | BioLegend | 104510 | 1:125 |
| Ly6G | PE-Cy7 | 1A8 | BioLegend | 127618 | 1:75 |
| CD64 | APC | X54-5/7.1 | BioLegend | 139306 | 1:100 |
| SiglecF | AF647 | E50-2440 | BD Biosciences | 562680 | 1:50 |
| Ly6C | AF700 | HK1.4 | BioLegend | 128024 | 1:100 |
| CD44 | APC-Fire750 | IM7 | BioLegend | 103062 | 1:100 |
| PD-1 | APC-Fire810 | 29F.1A12 | BioLegend | 135252 | 1:150 |
| F4/80 | BUV496 | T45-2342 | BD Biosciences | 75064 | 1:100 |
| CD80 | BUV661 | 16-10A1 | BD Biosciences | 741515 | 1:70 |
| CD3 | BUV805 | 17A2 | BD Biosciences | 741982 | 1:60 |
| Live/Dead | For V excitation | N/A[1] | Thermo Fisher | L34966 | 1:5000 |

[1]N/A: not applicable.
